# TREK-1 knock-down in human atrial fibroblasts leads to a myofibroblastic phenotype: a role in phenoconversion and overview of mechano-sensitive channel mRNA expression in cardiac diseases

**DOI:** 10.1101/2022.07.21.492231

**Authors:** Elisa Darkow, Dilmurat Yusuf, Sridharan Rajamani, Rolf Backofen, Peter Kohl, Ursula Ravens, Rémi Peyronnet

## Abstract

Cardiac cell mechanical environment changes on a beat-by-beat basis and also in the course of various cardiac diseases. Cells sense and adapt to such mechanical cues *via* specialized mechano-sensors mediating adaptive signaling cascades. Here, we report TREK-1 mRNA expression and activity in human atrial fibroblasts and reveal a cross talk between TREK-1 and fibroblast phenoconversion. With the aim to reveal additional candidates underlying mechano-transduction relevant to cardiac diseases, we investigated mechano-sensitive ion channels (MSC) mRNA expression in diseased and non-diseased human hearts.

Our results showed higher TREK-1 expression and activity in fibroblasts compared to myofibroblasts and we found that TREK-1 down-regulation leads a more myofibroblastic phenotype suggesting a role for this mechano-sensor in phenoconversion. In addition, TREK-1 is preferentially expressed in the left atrium compared to the right one and its expression is not significantly changed when fibroblasts from patients in sinus rhythm vs. sustained atrial fibrillation are compared. At the whole-heart level, numerous MSC were differentially expressed between atrial and ventricular or between non-diseased and diseased tissue samples.

Thus, we identify atrial fibroblast-specific TREK-1 expression and activity and reveal a role of TREK-1 in atrial fibroblast pheno-conversion. We also provide a comprehensive overview of cardiac MSC mRNA expression in atrial and ventricular tissue from diseased and non-diseased patients, identifying potential novel candidates underlying mechanotransduction in cardiac diseases.

## 1. Introduction

Cardiac cells are exposed to continually changing mechanical forces leading to stretch, compression, shear or a combination of these. Cardiac cells can sense, transduce and modify signaling pathways in response to mechanical cues. Molecules involved in the process of mechano-sensation and -transduction are integrins, various transmembrane receptors such as G-protein coupled receptors and receptor tyrosine kinases, and mechano-sensitive ion channels (MSC) [2].

The term MSC is here considered in its broadest sense, *i*.*e*. include mechanically modulated channels (MMC, channels responding indirectly to mechanical stimulation or requiring co-activation) and mechanically gated ion channels (MGC, channels responding directly to mechanical stimulation [Suppl. Figure 1]. MGC comprise stretch-activated ion channels (SAC) that activate in response to changes in membrane in-plane tension or curvature [3]. Based on their ion selectivity, SAC can be sub-divided into two main families: cation non-selective, and potassium (K^+^)-selective (SAC_NS_ and SAC_K_, respectively). The latter group includes the channel of our interest, *i*.*e*. TREK-1 (TWIK-related K^+^ channel, where TWIK stands for two-pore-domain weak inward rectifying K^+^ channel) [4, 5].

**Figure 1.**
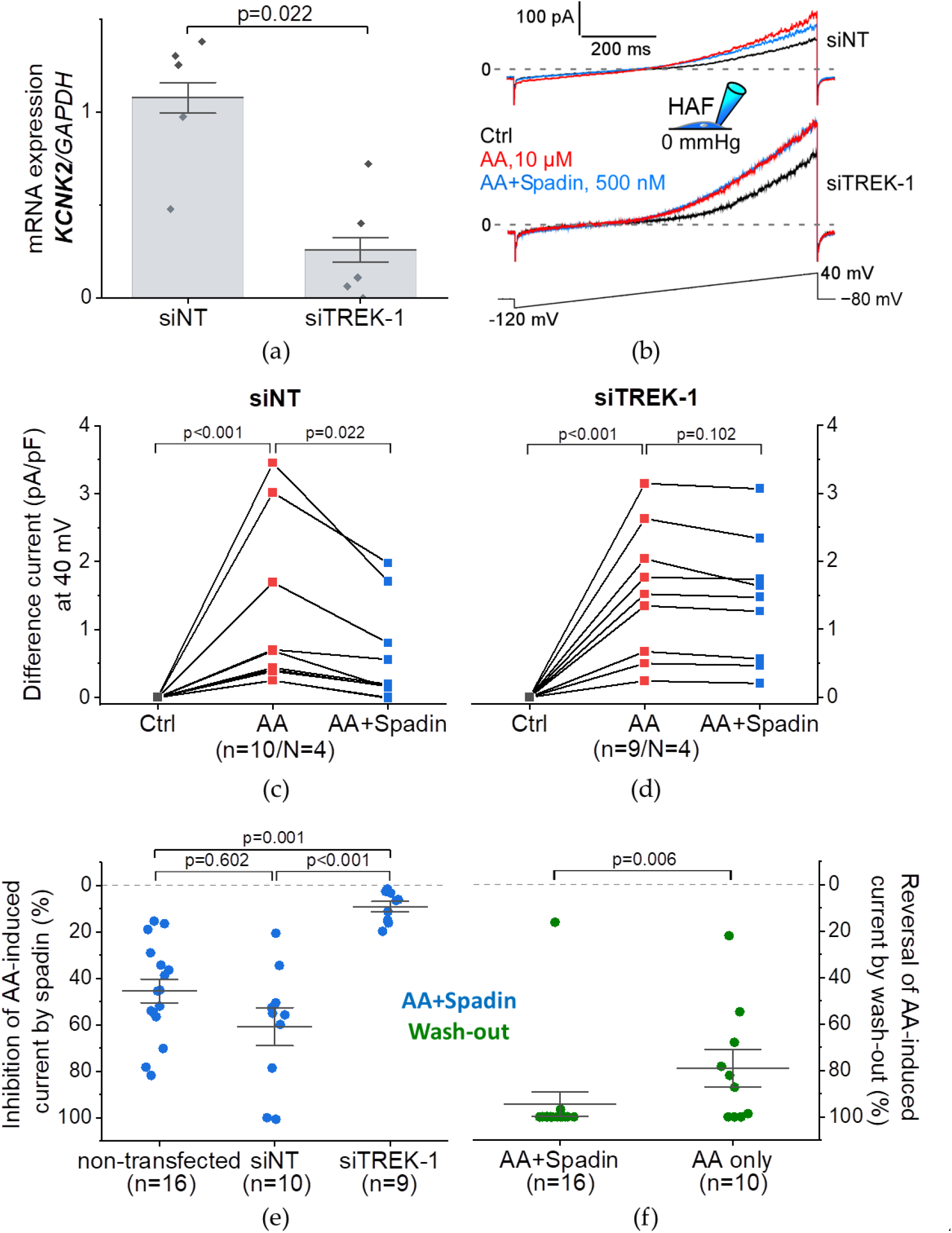
Arachidonic acid (AA)-induced and spadin-modulated currents in immortalized human atrial fibroblasts (HAF) without or with TREK-1 knock-down (siNT or siTREK-1, respectively). (a) qPCR results of *KCNK2* (TREK-1) mRNA expression normalized to the expression of *GAPDH* (housekeeping gene); 48 h after siRNA transfection; n = 5 dishes; N = 2 passages for both conditions; Mann−Whitney test. (b−f) Patch-clamp measurements in whole-cell configuration (pipette pressure 0 mmHg; b, inset) in response to a 800-ms-long ramp protocol from −120 to 40 mV (b, lowest trace). (b) Representative recording of absolute current with pre-drug control (Ctrl; black), in the presence of AA (10 µmol/L; red) and in the presence of AA and spadin (500 nmol/L; blue); top trace: si-TREK-1; middle trace: siNT. (c, d) Difference current density amplitudes (expressed in pA/pF) at 40 mV before (Ctrl; black), after exposure to AA (red), and after exposure to AA and spadin (blue) for siTREK-1 (*cf*. c) and siNT (*cf*. d); Paired samples Wilcoxon signed rank test. (e,f) Summary of the effect of spadin and wash-out on AA-induced currents in HAF. (e) Inhibition of AA-induced current by spadin (in %; blue) for cells without siRNA treatment (non-transfected, *cf*. **Error! Reference source not found**.), siNT (c) or siTREK-1 treatment (d); Kruskal−Wallis ANOVA with Dunn’s *posthoc* test. (f) Reversal of AA-induced current by wash-out (in %; green) for cells exposed to AA and spadin (AA+Spadin; *cf*. **Error! Reference source not found**.) or AA alone (*cf*. **Error! Reference source not found**.); Mann−Whitney test.

While mechano-transduction is important for physiological adaptive responses, excessive mechanical perturbation may lead to maladaptive tissue remodeling [6]. In this context, various transient receptor potential ion channels (TRP) that mediate stretch-induced calcium (Ca^2+^) influx have been associated with induction of fibrosis and arrhythmias [7–10], cardiac hypertrophy [11], heart failure [12] and myocardial infarction [13]. However, it remains unclear whether MSC remodeling occurs as a cause or a consequence of cardiac disease.

The SAC_K_ TREK-1 mediates background K^+^ currents. Being polymodal, it may change its conformation in response to a wide spectrum of stimuli, including temperature [14], membrane lipid composition [15], pH [16] and mechanical stress [15]. These ubiquitous stimuli are altered both under physiological and pathophysiological conditions. In the heart, TREK-1 is expressed in murine fibroblasts [17], but also in rat atrial [18, 19] and human ventricular cardiomyocytes [20]. In murine and human hearts, TREK-1 expression is higher in the ventricles than in the atria [21, 22] whereas in pigs, TREK-1 is predominantly expressed in the atria [23]. In a murine model, heart failure (HF) induced by transverse aortic constriction (TAC) was associated with upregulation of ventricular TREK-1 messenger ribonucleic acid (mRNA) [24]. TREK-1 downregulation in atrial fibrillation (AF) has been reported for right atrial (RA) protein expression in patients with lone AF [25] and for atrial mRNA expression in patients with concomitant HF [26]. In a patient with right ventricular outflow tract tachyarrhythmia, a point mutation in the selectivity filter of TREK-1 was shown to increase sodium (Na^+^) permeability and to induce hypersensitivity to stretch which may trigger arrhythmias [20]. Moreover, myocardial ischemia is connected to metabolic changes, such as extra- and intra-cellular acidosis [27], which in turn activate TREK-1. TREK-1 opening leads to K^+^ efflux and premature membrane repolarization, eventually promoting ventricular arrhythmias [28]. In a cell-type specific approach, TREK-1 knock-out in murine fibroblasts prevented cardiac dysfunction and reduced cardiac fibrosis in response to TAC [17]. Taken together, TREK-1 has been related to cardiac arrhythmogenesis and there are indications for a potentially central role for TREK-1 in cardiac fibroblasts in the context of fibrotic tissue remodeling but, to date, little is known about presence, relevance and function of TREK-1 in primary human atrial fibroblasts (HAF_prim_).

Molecular mechanisms underlying fibroblast mechano-sensing are not fully understood, but SAC have been proposed to play important roles [29, 30]. Besides being electrotonically coupled to cardiomyocytes and therefore being relevant to electrical conduction [31, 32], fibroblasts are the main producers of extra-cellular matrix (ECM) proteins, especially collagen type I and III. Under conditions of mechanical overload, cytokines like transforming growth factor (TGF)-β1 are secreted which leads to fibroblast “activation” into myofibroblasts, *i*.*e*. fibroblasts proliferate, differentiate into myofibroblasts (phenoconversion) and migrate [33, 34]. Compared with fibroblasts, myofibroblasts are larger in cell size and express substantial amounts of α-smooth muscle actin (αSMA) which confers contractile properties [35, 36]. Myofibroblasts also secrete elevated amounts of ECM proteins, contributing to tissue fibrosis and thereby causing tissue stiffening [37–41].

While mechano-regulation of various fibroblast functions may, at least initially, be part of a compensatory response (negative feedback), the resultant changes in tissue stiffness, sensed and transduced by fibroblasts’ SAC [42–44], may give rise to a vicious cycle (positive feedback) and contribute to fibrosis and the observation that “AF begets AF” [45]. Indeed, changes in tissue stiffness and mechanical overload accompany and promote AF [46–48]. Thus, several cardiac diseases, including AF itself, favor disease progression and maintenance by electrical, mechanical and structural remodeling [49]. For instance, in AF, mechanical stimuli such as atrial pressure or volume are increased [50], and fibrosis, a hallmark of AF, leads to profound changes in tissue structure, potentially forming obstacles to the propagation of the depolarization wave. Nevertheless, the mechanisms underlying domestication of AF remain unclear, and there are no causal treatment options targeting tissue remodeling in AF.

With this study, we aim to get a better understanding of the involvement of MSC in cardiac pathophysiology and to unravel possible candidates for chamber-selective drug therapy to avoid ventricular side effects. In particular, we aim to understand the roles of HAF_prim_ TREK-1 for AF-associated fibrotic tissue remodeling. We suggest that TREK-1 in HAF_prim_ acts as sensor and transducer of fibrotic remodeling.

Our result demonstrate that: i) TREK-1 is expressed and active in HAF_prim_ with a preferential expression in fibroblasts compared to myofibroblasts and in the left atrium compared to the right one, ii). TREK-1 is causally involved in the control of fibroblast phenoconversion and collagen gene expression, and iii) human cardiac MSC are differentially regulated (mRNA) between heart chambers and health and disease.

## 2. Materials and Methods

### 2.1. Cell culture

#### a) Cell types and maintenance

A cell line of immortalized HAF_prim_ (HAF) generated from the RA of a male patient in sinus rhythm (SR) [57] and 293T/17 human embryonic kidney (HEK) cells (CRL-11268, ATCC, Germany) were used for the sake of reproducibility and constant availability. Primary HAF_prim_ obtained by outgrowth and provided *via* the CardioVascular BioBank Freiburg, were used to check differences between RA and left atrium (LA) as well as SR and AF, and to account for inter-patient variability. For inter-species comparisons we also used pig primary RA and LA fibroblasts obtained by the outgrowth technique.

Fibroblast culture medium contained Dulbecco’s modified eagle medium [58], supplemented with GlutaMAX and sodium pyruvate (31966021, Thermo Fisher Scientific, Germany). HEK cell culture medium contained Dulbecco’s modified eagle medium (D6046, Sigma-Aldrich, Germany). Both media were supplemented with 10% (v/v) fetal bovine serum (F9665, Sigma-Aldrich) and 1% penicillin/streptomycin (P4333, Sigma-Aldrich). Fibroblasts and HEK cells were incubated at 37°C.

#### b) Cell passaging and seeding

For passaging of HAF and HEK cells at ≈90% confluency, the culture medium was discarded and cells were washed briefly with Dulbecco’s phosphate buffered saline (PBS; D1408, Sigma-Aldrich) [59]. Then, cells were detached using trypsin-ethylene diamine tetra-acetic acid (59418C, Sigma-Aldrich), transferred into 15 mL Falcon tubes and centrifuged (57 × g, 3 min, room temperature [20−22°C], Rotina 380, Hettich, Germany). The supernatant was discarded, the cell pellet re-suspended in culture medium and cells were re-seeded in 250 mL/75 cm^2^ cell culture flasks (658175, Greiner Bio-One, Germany) for subculture. For experiments, 30,000 – 60,000 cells were seeded *per* 2 mL/9.2 cm^2^ tissue culture dish (93040, Techno Plastic Products, Switzerland) or *per* 2 mL/9.6 cm^2^ well of a 6-well plate (657160, Greiner Bio-One). Cells were cultured for a minimum of 24 h to allow cell attachment before further protocols were applied.

#### c) Transfection

In HAF and HEK cells, TREK-1 was transiently knocked-down or overexpressed. Either transfection was performed 24 h after seeding.

For small interference RNA (siRNA)-based knock-down of TREK-1, HiPerfect reagent (301704, Qiagen, Germany) was used according to the manufacturer’s instructions and as described previously [44]. SMARTpool siRNA against human TREK-1 or as non-targeted control (L-006261-00-0005 and D-001810-10-05, respectively, Horizon, United Kingdom) were used at 8 nmol/L. Efficiency of the knock-down was assessed by quantitative real-time polymerase chain reaction (qPCR) 2 days post-transfection. Functional analyses were performed 3-4 days post-transfection.

For overexpression of human TREK-1 plasmid, jetPEI reagent (101-10N, Polyplus transfection, France) was used according to the manufacturer’s instructions and as described previously [60]. The TREK-1-internal ribosome entry site 2 (IRES2)-enhanced green fluorescent protein (eGFP) plasmid was kindly provided by Eric Honoré and Delphine Bichet. The IRES2-eGFP vector backbone alone was used as control. Cells were cultured for additional 4 days to allow expression of the protein before further experiments were performed. Successfully transfected cells were identified by cytosolic eGFP fluorescence.

#### d) Drug treatments

To obtain reference points, HAF phenoconversion was induced by well characterized compounds added to the cell culture medium. Treatment was initialized 24 h after seeding and lasted for 5-6 days.

Recombinant human TGF-β1 (0.4 nmol/L; ab50038, abcam, Germany) was added every second day to induce the myofibroblastic phenotype (solvent control: 0.1% double-distilled water). SD-208 (3 µmol/L; S7071, Sigma-Aldrich), a TGF-β receptor 1 kinase inhibitor, was renewed every day to induce the fibroblastic phenotype [36, 61, 62] (solvent control: 0.03% dimethyl sulfoxide [DMSO; D8418, Sigma-Aldrich]).

For characterization of SD-208 and TGF-β1 treatment, reversibility and the time course of fibroblast phenoconversion were tested. The protocol to test for the reversibility extended to 13 days, with switchover of the treatment on day 7 (after 6 days of treatment). QPCR samples for time course analyses were collected daily during the 6 day treatment protocol. Morphological changes during drug treatment were monitored with an inverted microscope (Eclipse TS100, Nikon, Japan).

#### e) Outgrowth

HAF_prim_ were isolated using the outgrowth method as described earlier [63]. Conceptually, fibroblasts grow out of myocardial chunks placed into culture medium due to their migratory and proliferative potential.

To this end, human RA and/or LA tissue was received from patients in SR or AF undergoing open-heart surgery at the Universitåts-Herzzentrum Freiburg· Bad-Krozingen *via* the CardioVascular BioBank Freiburg. Patients gave written informed consent to the use of the tissue (with approval by the Ethics Commission of the University of Freiburg, reference: 393/16: 214/18). Patients were stratified according to their pre-surgical rhythm status. Here, samples from patients in SR were taken to reflect control properties with regard to the heart rhythm, even though they were not obtained from healthy patients. In contrast, samples from patients in sustained AF were expected to exhibit profound electrical and structural alterations. Due to higher availability of tissue from patients in sustained AF than in paroxysmal AF, only data for sustained AF were included in this study and will be referred to as AF subsequently. Clinical characteristics of the patients whose tissue was used for outgrowth culture are summarized in **Table 1**.

**Table 1.**
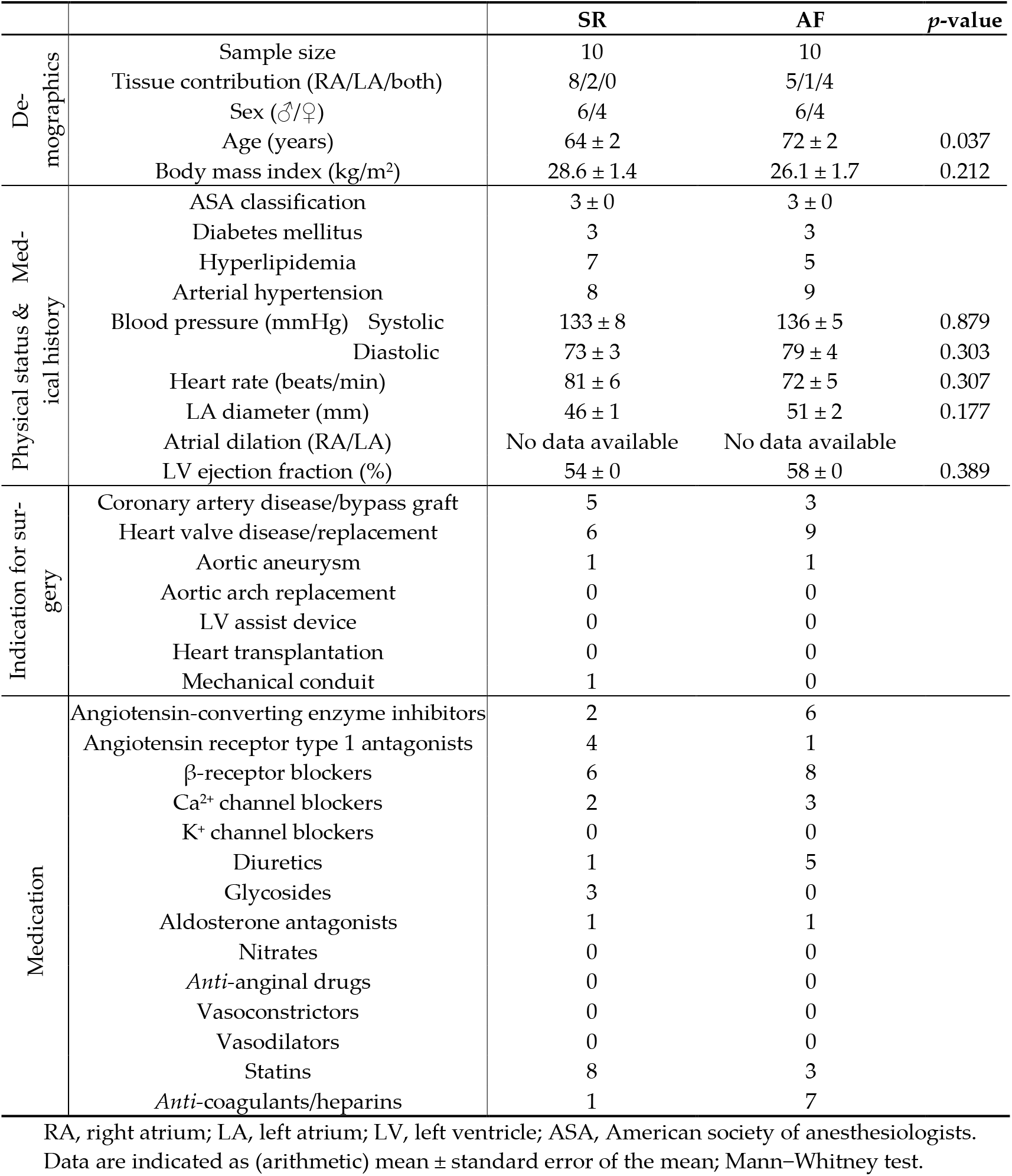
Clinical data of patients whose tissue provided the HAF_prim_ used in this study. Demographics, physical status, medical history, indication for surgery and long-term medication before surgeries were compared between patients with sinus rhythm (SR) and patients with sustained atrial fibrillation (AF).

For outgrowth, 1-5 atrial tissue chunks of ≈1 mm^3^ were placed into 2 mL/9.2 cm^2^ tissue culture dishes (93040, Techno Plastic Products) with scarped surfaces. Then, the chunks were covered with 2 mL fibroblast cell culture medium which was exchanged twice a week. After 21 days of culture and without passaging, the HAF_prim_ were used for assessment of TREK-1 and αSMA mRNA expression, and of stretch-induced activity by qPCR and patch-clamp, respectively.

The same protocol was applied for pig RA and LA fibroblasts. Ethical approval for *ex vivo* pig heart experiments has been obtained in accordance with German ethical guidelines for laboratory animals and approved by the Institutional Animal Care and Use Committee of the City of Freiburg and the University of Freiburg (authorization: X-21/03B).

### 2.2. qPCR

We used qPCR to assess mRNA expression levels of TREK-1 (*KCNK2*), αSMA (*ACTA2*), collagen type I α-1 (*COL1A1*), collagen type I α-2 (*COL1A2*), collagen type III α-1 (*COL3A1*), Piezo1 (*PIEZO1*), Piezo2 (*PIEZO2*), and α-1 subunit of Ca^2+^-activated K^+^ channel of large (“big”) conductance (BK_Ca_ α; *KCNMA1*). These mRNA expression levels were normalized to the mRNA expression level of glyceraldehyde-3-phosphate dehydrogenase (GAPDH; *GAPDH*).

Isolation of fibroblast mRNA was performed with a commercial RNA isolation kit (74004, Qiagen) following the manufacturer’s instructions. Briefly, fibroblasts were washed once with PBS (containing in mmol/L: 137 NaCl, 2.7 KCl, 10 Na_2_HPO_4_, 1.8 KH_2_PO_4_, pH 7.4 adjusted with HCl, 300±10 mOsm/L [osmometer: K-7400, Knauer, Germany]) and lysed in 350 µL RLT lysis buffer (79216, Qiagen) *per* dish/well supplemented with 1% (v/v) β-mercaptoethanol (1610710, Bio-Rad, Germany). Fibroblasts were then scraped off the plates (32 702180, SLG, Germany), transferred into 1.5 mL tubes and stored at −80°C until further processing. After complete mRNA isolation including multiple centrifugation steps (centrifuge 5424 R, Eppendorf, Germany), total mRNA concentration was determined photospectrometrically (ND-1000, Thermo Fisher Scientific). 100 ng mRNA/sample were converted into complementary deoxyribonucleic acid (cDNA) using reverse transcription reagents (N8080234, TaqMan, Germany). CDNA was amplified in qPCR master mix (4444556, TaqMan) during 45 cycles (LightCycler 480 System, Roche Diagnostics, Switzerland) using the assays Hs02786624_g1 for *GAPDH*, Hs01005159_m1 for *KCNK2*, Hs00426835_g1 for *ACTA2*, Hs00164004_m1 for *COL1A1*, Hs01028956_m1 for *COL1A2*, Hs00943809_m1 for *COL3A1*, Hs00207230_m1 for *PIEZO1*, Hs00401026_m1 for *PIEZO2*, and Hs01119504_m1 for *KCNMA1* (all TaqMan).

### 2.3. Patch-clamp

To record ion channel activity in the plasma membrane, the patch-clamp technique in voltage-clamp mode was used as described previously [60]. Recordings were obtained in whole-cell (electrical stimulation) or inside-out (mechanical stimulation) configurations. The presence of TREK-1 activity was assessed by the voltage dependence and stretch-induction of the current, and by pharmacological probes.

All measurements were performed at room temperature and cell culture dishes were exchanged every hour. To test for TREK-1 (reversal potential E_rev,K_ ≈−80 mV), voltage was clamped to 0 mV, largely excluding SAC_NS_ (E_rev,NS_ ≈ 0 mV). Seal resistances accepted were ≈1 GΩ, independent of pipette size and cell quality. Currents were acquired at 20 kHz sampling rate, and low-pass filtered at 1 kHz. Recordings were acquired with pCLAMP 10.6 and analyzed with Clampfit 10.6 (both softwares, Molecular Devices, United States). Representative recordings were selected to represent average current amplitude and morphology. Liquid junction potential are not corrected.

#### a) Whole-cell configuration

For whole-cell configuration, pipette solution, formulated to mimic the intra-cellular ionic environment, contained (in mmol/L): 155 KCl, 3 MgCl_2_, 5 EGTA, 10 N-2-hydroxy-ethylpiperazine-N-ethanesulfonic acid (HEPES), pH buffered to 7.2 using KOH, ≈300 mOsm/L. The solution was stored at room temperature. Bath solution, formulated to mimic the extra-cellular ionic environment, contained (in mmol/L): 150 NaCl, 5 KCl, 10 HEPES, 2 CaCl_2_, pH buffered to 7.4 using NaOH, ≈300 mOsm/L. The solution was prepared ahead of experiments and stored in aliquots at −20°C.

Average pipette resistance was 1.88 ± 0.02 MΩ, n = 45. For electrical stimulation, a voltage ramp protocol with continuous change from −120 mV to 40 mV within 0.8 s was utilized. Between application of ramps, cells were held at −80 mV for 5 s.

A local perfusion system was installed for controlled administration of bath solution (control), or one of the following compounds: arachidonic acid (AA, 10 μmol/L; SML1395, Sigma-Aldrich), a non-selective TREK-1 activator [64, 65], or spadin (500 nmol/L; 5594, Tocris, Germany), a TREK-1-selective blocker [66, 67]. The effect of simultaneous perfusion of AA and spadin had been verified before, showing that spadin selectively antagonizes the action of AA [67].

During data analysis, currents were normalized to membrane capacitance (a surrogate for cell size) and expressed as current density (in pA/pF). Currents at the maximum level of depolarization (40 mV) were analyzed for each ramp acquired.

#### b) Inside-out configuration

For inside-out recordings, the cell-attached configuration was established and then, the patch pipette was rapidly retracted from the cell. Indeed, TREK-1 is inhibited by the actin cytoskeleton [68], which is disrupted in inside-out configuration. Pipette and bath solutions were inverted, compared to whole-cell recordings. In order to dissect TREK-1 activity from that of other K^+^ conducting channels (BK_Ca_, K_v_, or K_ATP_), we added established inhibitors to the pipette solution: 3 mmol/L 4-aminopyridine (4-AP; A78403), 10 mmol/L tetraethylamonium (TEA; T2265), 10 µmol/L glibenclamide (G0639, all Sigma-Aldrich) [69]. These inhibitors do not suppress K_2P_ channel activity [69, 70].

Average pipette resistance was 1.38 ± 0.01 MΩ, n = 407. For mechanical stimulation, negative pressure was applied for 500 ms, in 10 mmHg increments from 0 to −80 mmHg (9 sweeps), with 1.8 s at 0 mmHg between pressure steps, using an automated precision system (HSPC-1, ALA Scientific Instruments, United States).

During data analysis, baseline activity at 0 mmHg was taken as 0 pA reference (average baseline current = −1.17 ± 0.04 pA). Peak, average and near steady-state currents were analyzed during pressure pulses (illustrated in **Error! Reference source not found**. b). Single channel activity was also analyzed during pressure pulses. Recordings of silent cells were defined as flat traces without deviations from the baseline current that would have been characteristic for ion channel activity. Active cells were defined as having traces with >1 pA current spikes that show characteristic ion channel kinetics (single channel openings and closures are clearly distinguishable). Among the active cells, we discriminated four current patterns. First, TREK-1-like currents were characterized by a single channel amplitude ≈2 pA (established in **Error! Reference source not found**. a,b), characteristic kinetics, and long open dwell times (≈50 ms) [15]. Second, BK_Ca_-like currents were characterized by a single channel amplitude ≈5 pA, characteristic kinetics, and shorter open dwell times (≈20 ms) [63]. Third, Piezo1-like currents were characterized by an accumulation of single channel amplitudes (≈1 pA each), without characteristic kinetics and with temporarily overlapping channel activities [63]. The fourth current pattern was characterized by a single channel amplitude ≈1 pA, without characteristic kinetics and without temporarily overlapping channel activities. One cell could have displayed multiple current patterns.

### 2.4. Imaging

#### a) Immunocytochemistry

For imaging of αSMA, a marker of myofibroblasts, HAF were seeded onto Ø 10 or 16 mm borosilicate glass cover glasses (631-0170 and 631-0151, VWR, Germany). After transfer into 12- or 24-well plates (665102 and 662160, respectively, Greiner Bio-One), cells were chemically fixed with 4% para-formaldehyde solution. For staining of αSMA, mouse anti- αSMA primary antibody (A5228, Sigma-Aldrich) and goat anti-mouse Alexa Fluor 488 secondary antibody (1:500, 1h; A-21121, Invitrogen, Germany) were used. Nuclei were counterstained using 4′,6-diamidin-2-phenylindol (DAPI, 1:1000, 7 min; D1306, Thermo Fisher Scientific). Washing steps were performed with PBS. Coverslips were mounted using PermaFluor aqueous mounting medium (12695925, Thermo Fisher Scientific).

#### b) Confocal microscopy

Imaging was performed using an inverted confocal microscope (TCS SP8 X, Leica Microsystems, Germany) at the microscopy facility SCI-MED (Super-Resolution Confocal/Multiphoton Imaging for Multiparametric Experimental Designs, Institute for Experimental Cardiovascular Medicine, Freiburg, Germany). Image acquisition was performed with the Application Suite X software (Leica Microsystems). All measurements were conducted in PBS at room temperature.

To get an overview, a 40× water objective or a dry 10× objective were used. For final image acquisition, a 63× oil objective (numerical aperture = 1.4) was used. The 488 nm line of a white light laser was used to excite Alexa Fluor 488; a 405 nm diode laser was used to excite DAPI nucleus staining. The focal plane was set at the maximal cell cross-section area.

Cells’ cross sectional areas and fluorescence intensities analyses were performed with ImageJ software [71]. Only cells that appeared viable, *i.e*. showing a sharp and contrasted plasma membrane, a well-delineated nucleus, and cell attachment to the dish (presence of membrane protrusions) were considered. Cell borders were manually outlined based on the transmitted light image.

### 2.5. Affymetrix

Gene expression in HAF_prim_ from patients with SR or AF was assessed by *de novo* analysis of the gene micro-array dataset first published by Poulet *et al*. in 2016 [72]. Briefly, cells were isolated using the outgrowth technique, cultured for 21 days, passaged once and cultured for another 10-15 days. Isolated RNA was processed for GeneChip Human Gene 1.0 ST Array (Affymetrix, United States) according to the manufacturer’s instructions.

### 2.6. RNA-Sequencing

#### a) Studies considered for the meta-analysis

National center for Biotechnology Information (NCBI) sequence read archive (SRA) database [73] allowed to access deposited transcriptomic datasets. Studies had to fulfill the following criteria for inclusion into this meta-analysis: homo sapiens (organism), cardiac (tissue type), RNA (source), RNA-sequencing (strategy), Illumina (platform), paired-end (library layout), fastq (file type). The sequencing files of human cardiac tissue samples were then further selected based on the patients’ health status: non-diseased (no structural heart disease), dilated cardiomyopathy (DCM), ischemic cardiomyopathy (ICM), coronary artery disease (CAD), heart valve disease (HVD) or AF. Non-diseased atrial and ventricular myocardial samples originated from donor hearts not suitable for transplantation due to size mismatch or logistic reasons. Diseased myocardial samples were obtained from patients undergoing open heart surgery. All patients gave informed consent for the respective study. Only tissue from patients with sustained (and not paroxysmal) AF was included in the AF group. Of note, all patients with AF also had CAD or HVD as reason for cardiac surgery but tissue samples were only grouped into AF. Furthermore, the samples had to originate from (tissue provenance) the left ventricle (LV), right ventricle (RV), LA or RA. This meta-analysis included paired (different provenances from the same individual) and non-paired tissue samples. In order to relate better our results to published work, especially results of functional experiments, LV samples with DCM and ICM constitute the HF group and samples from RA with CAD and HVD compose the SR group. Considering these inclusion criteria, the analysis of RNA-sequencing studies, with a total of 108 samples from 62 individuals, is summarized in **Table 2**. Details on sample acquisition, including ethical approvals, and technical procedure can be found in the corresponding original publications. Patient characteristics are summarized in **Table 3**.

**Table 2.**
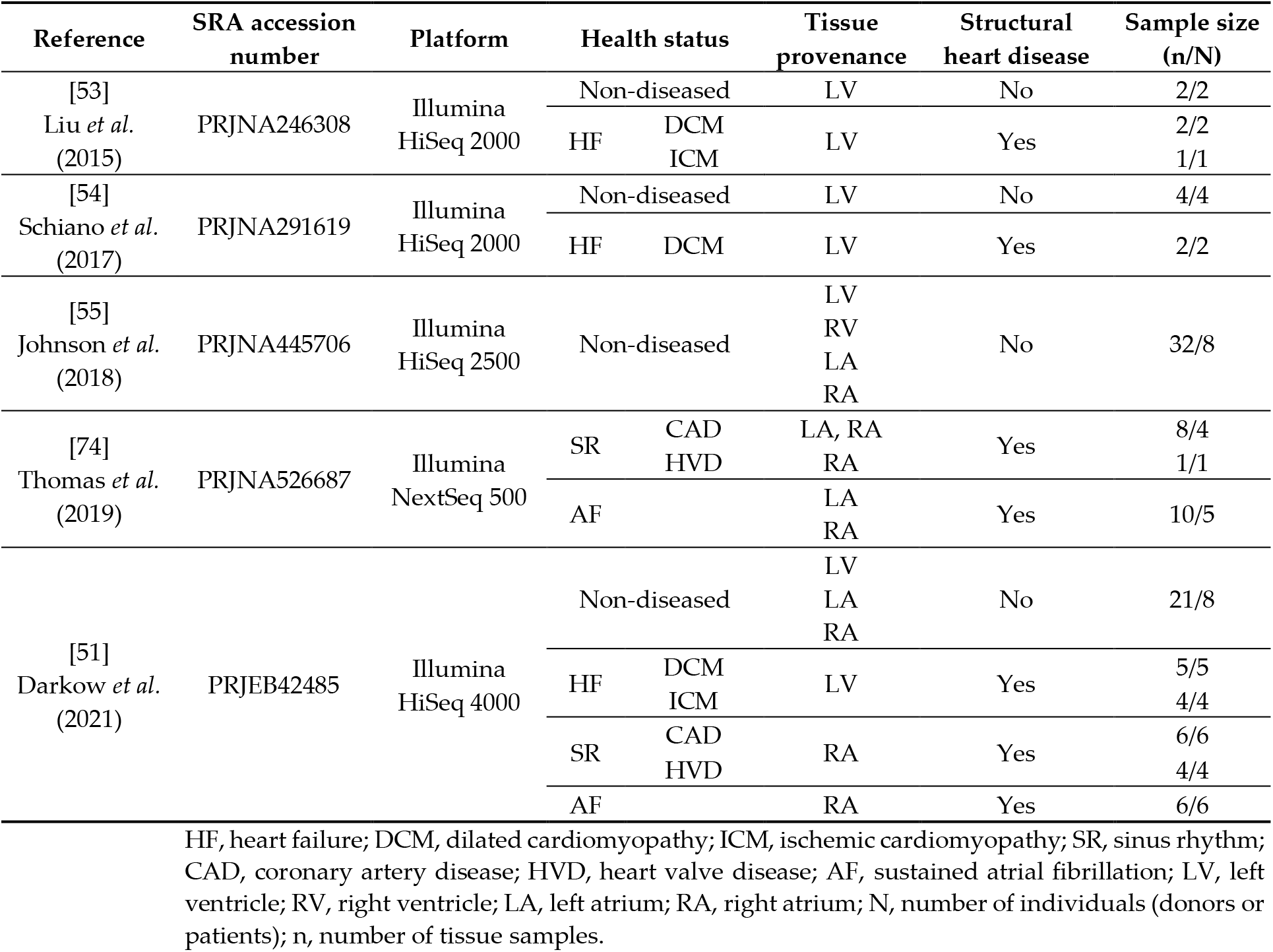
Studies considered for the meta-analysis.

**Table 3.**
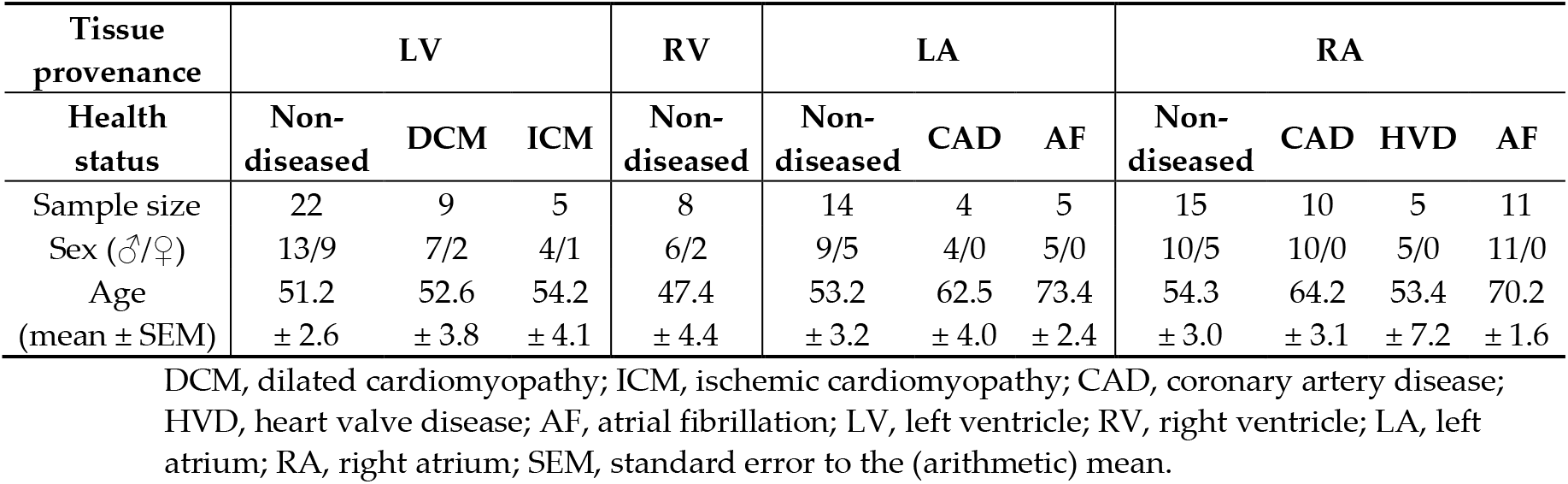
Clinical data of patients included in the RNA-sequencing meta-analysis.

| Tissue provenance | LV |  |  | RV | LA |  |  | RA |  |  |  |
| --- | --- | --- | --- | --- | --- | --- | --- | --- | --- | --- | --- |
| Health status | Non-diseased | DCM | ICM | Non-diseased | Non-diseased | CAD | AF | Non-diseased | CAD | HVD | AF |
| Sample size | 22 | 9 | 5 | 8 | 14 | 4 | 5 | 15 | 10 | 5 | 11 |
| Sex (♂/♀) | 13/9 | 7/2 | 4/1 | 6/2 | 9/5 | 4/0 | 5/0 | 10/5 | 10/0 | 5/0 | 11/0 |
| Age | 51.2 | 52.6 | 54.2 | 47.4 | 53.2 | 62.5 | 73.4 | 54.3 | 64.2 | 53.4 | 70.2 |
| (mean ± SEM) | ± 2.6 | ± 3.8 | ± 4.1 | ± 4.4 | ± 3.2 | ± 4.0 | ± 2.4 | ± 3.0 | ± 3.1 | ± 7.2 | ± 1.6 |

DCM, dilated cardiomyopathy; ICM, ischemic cardiomyopathy; CAD, coronary artery disease; HVD, heart valve disease; AF, atrial fibrillation; LV, left ventricle; RV, right ventricle; LA, left atrium; RA, right atrium; SEM, standard error to the (arithmetic) mean.

#### b) Data Analysis

The RNA-sequencing data analysis was mainly carried out using the Galaxy platform [75], following guidelines from the tutorial “Reference-based RNA-Seq data analysis” [76, 77]. In short, data were downloaded and extracted in fastq format from NCBI SRA [73]. Thomas *et al*. deposited 20 runs/sample to achieve higher coverage. All runs for one sample, *i.e*. fastq files separated into forward and reverse datasets, were merged using Concatenate datasets tail-to-head [78]. Quality control checks on raw sequence data were performed using FastQC [79] and MultiQC [80]. Then, Cutadapt [81] was used to remove adapter sequences. The splice-aware aligner STAR [82] was used to map the RNA-sequencing reads onto the human reference genome (hg19). The mapping results were visualized by the integrative genome viewer, IGV [83]. Thereafter, the strandness of the RNA-sequencing data (reads mapping to the forward or reverse DNA strand) was estimated using Infer Experiment from RSeQC [84]. Gene expression was measured by featureCounts [85].

From the published reference datasets, we carefully selected and combined the samples into three different groups of pairwise comparisons where the condition samples were balanced in each respective group: AF *vs*. SR (from RA tissue samples), HF *vs*. nondiseased (from LV tissue samples), and atria *vs*. ventricles (from non-diseased tissue samples). For the comparison of AF with SR, we selected 25 samples from the reference datasets of Thomas *et al*. [74] and Darkow *et al*. [51]. To compare HF with non-diseased, we selected 28 samples from the reference datasets of Darkow *et al*., Schiano *et al*. [54] and Liu *et al*. [53]. To compare atria with ventricles, we selected 53 samples from the reference datasets of Johnson *et al*. [55] and Darkow *et al*. For each pairwise comparison, we used the DESeq2 package [86] in R [87] to test for differential expression of the genes, and employed the gglot2 package [88] to visualize the overall effect of experimental covariates with the plots of principal component analysis (PCA). When running DESeq2 function on raw counts, to control for the batch effects derived from the different references, we adopted the model design ∼ batch + condition, where batch corresponds to the reference, and condition to the health status or the tissue provenance. The resulting “normalized” counts were transformed by the function vst() of DESeq2 and further underwent batch removal with the function removeBatchEffect() of limma package [89], which served as the input to generate the PCA plots. For checking differential expression, we first ran the function DESeq(), and then applied the function lfcShrink() with the shrinkage estimator ashr [90]. Differential gene expression levels with adjusted *p*-values < 0.05 (Benjamini-Hochberg procedure) were regarded as significantly different. DESeq2 output files were re-uploaded to Galaxy platform and used to generate the heatmap and the volcano plots [88].

### 2.7. Statistical Analysis

Unless otherwise indicated, values are expressed as arithmetic mean ± standard error of the mean. In the following, arithmetic mean is referred to as average. For HAF, N-numbers refer to the number of passages. For primary fibroblasts, N-numbers refer to the number of tissue samples, *i.e*. number of patients. In both cases, n-numbers refer to the number of individual cells/dishes.

Descriptive statistics (arithmetic mean, geometric mean, median, skewness, curtosis) and Shapiro-Wilk normality test were used to assess the distribution of data. After graphical evaluation using a scatter plot, we tested for outliers with Grubb’s test, if needed.

Significance of the difference between averages of two conditions was assessed using the unpaired t-test (parametric), the Wilcoxon signed rank test (non-parametric, paired) or the Mann−Whitney test (non-parametric, unpaired). In case of unpaired t-test, the homogeneity of variances was tested with the F-test. Significance of the difference between averages of more than two conditions was assessed using the non-parametric Kruskal−Wallis (unpaired) analysis of variance (ANOVA) with Dunn’s (all with all) or Dunnett’s (all to reference) *post-hoc* test.

Statistical analysis and data representation were performed with OriginPro 2020 (United States). Dunnett’s *post-hoc* test was performed with R package DescTools [87, 91]. ns indicates non-significant; * indicates *p* ≤ 0.05; ** indicates *p* < 0.01; *** indicates *p* < 0.001.

## 3. Results

### 3.1. TREK-1 is expressed and active in HAF

In this study, we first assessed TREK-1 expression and activity in the immortalized cell line HAF. On average, siRNA-based TREK-1 knock-down (siTREK-1) reduced *KCNK2* (TREK-1) mRNA expression by 76% compared to non-targeted siRNA treatment (siNT, control; **Figure 1** a). In these two conditions, we investigated TREK-1 activity in wholecell patch-clamp configuration while superfusing the non-selective TREK-1 activator, AA (10 µmol/L), in combination with the selective TREK-1 inhibitor, spadin (500 nmol/L). A ramp protocol from −120 to 40 mV revealed an outward rectifying background current in siTREK-1 and siNT HAF. Exposure of these cells to AA increased current amplitudes (**Figure 1** b−d, in red). AA-induced activation increased over time and reached significance from 30 s onwards. This response was reversible upon wash-out of AA (**Error! Reference source not found**.).

In order to assess whether TREK-1 channels contribute to the AA-sensitive current in HAF, the selective TREK-1 blocker, spadin, was added on top of AA after 80 s. This gave rise to a decrease in current amplitude in siNT cells, but not in siTREK-1 cells (**Figure 1** b−d, in blue). Paired difference current densities (at 40 mV; normalized to pre-drug control) from 10 siNT cells showed that AA increased current density by 1.1 ± 0.4 pA/pF; this effect was reversed upon additional exposure to spadin (average current density decrease −0.6 ± 0.2 pA/pF; **Figure 1** c). Same results were observed in paired data from 16 non siRNA-transfected HAF (**Error! Reference source not found**.). In 9 siTREK-1 cells, spadin did not reverse the current increase induced by AA (average current density Δ −0.1 ± 0.3 pA/pF; **Figure 1** d).

In order to account for the large variability of responses to AA, the inhibitory effects of spadin were additionally expressed in percent of AA-induced current. Spadin reduced AA-activated current in non-transfected HAF by 45.4 ± 5.1 % and by 60.8 ± 8.2 % in siNT transfected cells, but had no significant effect in siTREK-1 cells (9.1 ± 2.1%) (**Figure 1** e). The AA-induced current activation was largely reversible upon wash-out either of AA and spadin (94.6 ± 5.2%) or of AA alone (83.8 ± 10.1%) (**Figure 1** f). This combination of approaches confirmed TREK-1 expression and activity in HAF.

Importantly, the current density upon exposure to AA was not significantly different between siTREK-1 and siNT treated HAF (*cf*. **Figure 1** c, d; *p* = 0.348 [Mann−Whitney test]), highlighting the contribution of other AA-sensitive ion channels. Indeed, micro-array gene expression data from HAF_prim_ of patients with SR showed mRNA expression of various AA-sensitive ion channels (**Error! Reference source not found**. a). Among them, *KCNMA1* (BK_Ca_ subunit α 1) was robustly expressed, matching with the large single channel events observed at depolarized potentials (around 40 mV) and in recordings using the inside-out configuration (**Error! Reference source not found**. b, c).

### 3.2. TREK-1 is higher expressed and active in fibroblasts than in myofibroblasts

We hypothesized that the high inter-cellular variability observed in **Figure 1** has been due to different fibroblast phenotypes in the HAF cell line. To assess this hypothesis, HAF were differentiated into myofibroblasts by exposure to TGF-β1 (400 pmol/L, solvent control: 0.1% water), and into fibroblasts by exposure to the selective TGF-β receptor 1 kinase inhibitor SD-208 (3 µmol/L, solvent control: 0.03% DMSO). Taken together, the results suggested that HAF are pre-activated (no significant effect of TGF-β1) while SD-208 robustly reduced the percentage of αSMA positive cells, cell area, and mRNA expression of *COL1A1, COL1A2* (collagen type I) and *ACTA2* (αSMA; **Error! Reference source not found**. and **Suppl. Error! Reference source not found**.).

*KCNK2* mRNA expression was examined in the different fibroblast phenotypes. *KCNK2* mRNA expression was upregulated under SD-208 treatment in comparison to TGF-β1 and DMSO (2.0 ± 0.6 *vs*. 0.2 ± 0.0 and 0.3 ± 0.1, respectively; **Figure 2** a). A similar effect was observed on *KCNMA1* mRNA expression, *i.e*. upregulation under SD-208 treatment in comparison to DMSO and medium only cultured HAF. *PIEZO1* and *PIEZO2* (Piezo1,2; both SAC_NS_) showed no significantly altered gene expression with either treatment (**Error! Reference source not found**.).

**Figure 2.**
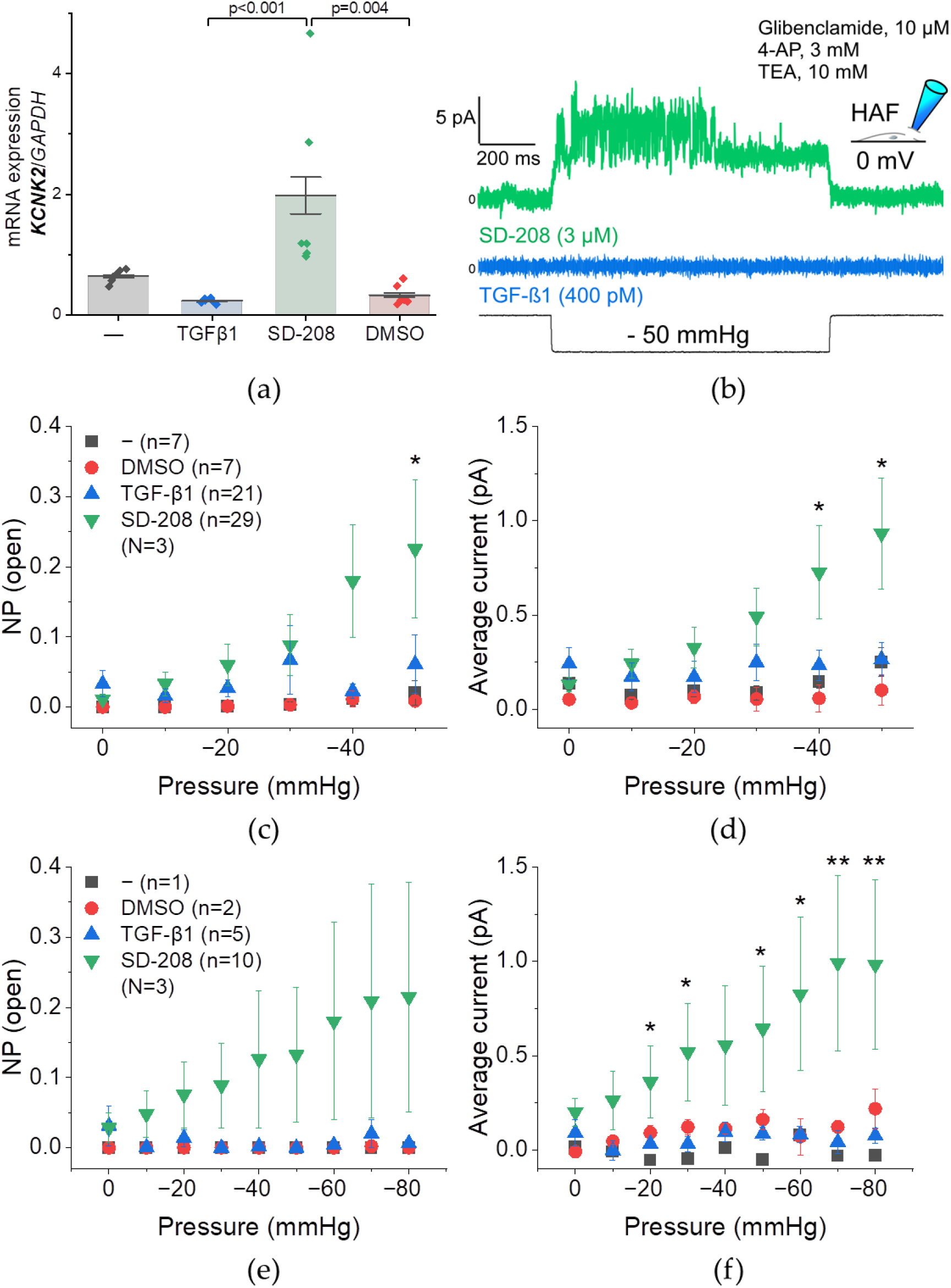
TREK-1 mRNA expression and stretch-induced activity in HAF. Cell culture medium only (−), or supplemented with TGF-β1 (400 pmol/L, blue) or SD-208 (3 µmol/L, green; solvent control: 0.03% DMSO, red) were compared. (a) *KCNK2* (TREK-1) mRNA expression determined by qPCR and normalized to the expression of *GAPDH* (housekeeping gene); n = 6 wells/N = 3 passages for all conditions; Kruskal−Wallis ANOVA with Dunn’s *post-hoc* test (only significant differences are indicated). (b−f) Patch-clamp measurements in inside-out configuration at a holding potential of 0 mV in the presence of non-K_2P_ channel inhibitors (b, inset). (b) Representative recording of a SD-208 treated cell (green) and a TGF-β1 treated cell (blue) during application of negative pressure to the patch of −50 mmHg; electrical interference (50 Hz) and Notch center (50 Hz) filters applied. (c, e) Open probability−pressure relationship for 2 pA single channel amplitudes. (d, f) Average current (over the duration of the pressure pulse)−pressure relationship. Only cells that sustained −50 mmHg (c, d) or −80 mmHg (e, f) were included; n = number of cells; N = number of passages; * indicates significance between SD-208 and TGF-β1 assessed by unpaired t-test (c, d) or Mann−Whitney test (e, f).

Next, we examined whether the higher mRNA expression of *KCNK2* in SD-208 treated fibroblasts translated into higher TREK-1 activity in the plasma membrane compared to TGF-β1 treated myofibroblasts. To better distinguish TREK-1 activity compared to the initial characterization (*cf*. **Figure 1**), K^+^ channel inhibitors which do not affect K_2P_ channel activity were added to the pipette solution. In order to see more easily TREK-1 activity, the inside-out patch configuration with mechanical stimulation was used (TREK-1 is inhibited by the cytoskeleton, thus more active in inside-out as the cytoskeleton is desorganised; protocol controlled in HEK cells transiently overexpressing TREK-1; **Error! Reference source not found**.). Single channel events of ≈ 2pA amplitude were analyzed as this matches TREK-1 features when overexpressed in HEK cells (**Error! Reference source not found**.).

We found that the activity in fibroblasts had higher open probability at −50 mmHg (0.23 ± 0.10 *vs*. 0.06 ±0.04) and higher average current at −40 and −50 mmHg (at −50 mmHg: 0.93 ± 0.29 pA *vs*. 0.26 ± 0.09 pA) when considering cells sustaining −50 mmHg compared to myofibroblasts (**Figure 2** b−d). In few cells, the entire stretch protocol (with suction levels up to −80 mmHg) could be recorded. TREK-1 activity was higher in fibroblasts than in myofibroblasts for −20 mmHg and more negative pressure pulses (excluding −40 mmHg; at −80 mmHg: 0.98 ± 0.45 pA *vs*. 0.08 ± 0.04 pA) but no significant differences in open probability were seen, mainly due to the high variability (**Figure 2** e, f).

To further characterizethe interrelation between *KCNK2* expression and fibroblast phenoconversion, we investigated the reversibility (**Figure 3** a) and the time course (**Figure 3** b) of HAF phenoconversion with SD-208 and TGF-β1.

**Figure 3.**
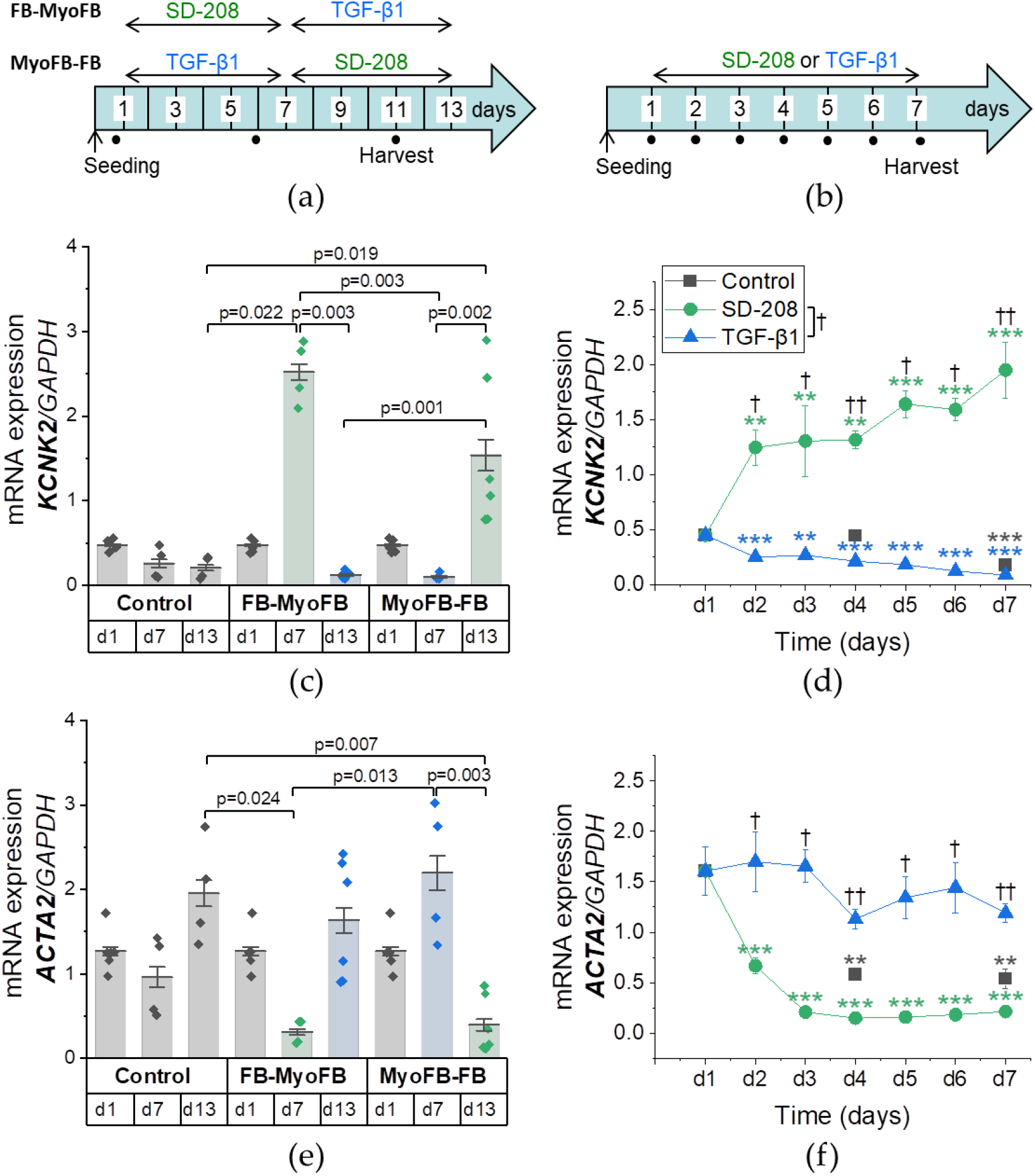
Reversibility and time course of fibroblast (FB)-myofibroblast (MyoFB) differentiation in HAF. qPCR results for *ACTA2* (αSMA) and *KCNK2* (TREK-1) mRNA expression normalized to *GAPDH*. (a, b) Seeding into cell culture dishes at d0; treatment from d1 to d7 (b) or d13 (a) with TGF-β1 (400 pmol/L, blue) or SD-208 (3 µmol/L green; solvent: DMSO); control: medium exchange only (grey). (c, e) Phenotype reversibility; d7: n = 2 dishes/condition; day 13: n = 3 dishes/condition; N = 2 passages; Kruskal−Wallis ANOVA with Dunn’s *post-hoc* test (only significant differences are indicated). (d, f) Phenoconversion time course; d1: the same three control data points are plotted for each condition; n = 2−3 dishes/time point; N = 2 passages; significance: * for comparison of different time points to d1 for the same treatment (Kruskal−Wallis ANOVA with Dunnett’s *post-hoc* test) and † for comparison of different treatments for the same time point (Mann−Whitney test on d2, d3, d5, d6 and Kruskal−Wallis ANOVA with Dunn’s *post-hoc* test on d4, d7); On d4 and d7 no significant difference between the control and SD-208 and/or TGF-β1 was detected.

*KCNK2* mRNA expression was modulated in the opposite direction than *ACTA2* mRNA expression and was reversible upon SD-208 or TGF-β1 treatment (**Figure 3** c, d). SD-208 and TGF-β1 treatment resulted in an increase or decrease, respectively, in *KCNK2* mRNA expression within 1 day of exposure (**Figure 3** d). SD-208 reversed myofibroblasts into fibroblasts, *i.e*. in cells pre-treated with TGF-β1 we observed a decrease in *ACTA2* mRNA expression within 1 day which stayed reduced throughout the 6 days of SD-208 treatment (**Figure 3** e). TGF-β1 treated cells retained *ACTA2* expression suggesting a high pre-activation of the cell line (**Figure 3** f), as previously highlighted.

*KCNK2* mRNA expression decreased over time in control conditions (medium exchange only; **Figure 3** d). *ACTA2* mRNA expression in control conditions showed that this cell line has an inherent tendency to become more fibroblastic between day 1 and day 7 of culture (**Figure 3** f). After 13 days, HAF pre-treated with TGF-β1 detached upon subsequent SD-208 treatment (≈ 15% confluency), but HAF pre-treated with SD-208 did not detach upon subsequent TGF-β1 treatment or medium exchange only (≈ 100% confluency; data not shown).

Taken together, we observed that SD-208 and TGF-β1 took effect after 1 day of incubation. These modulators counteracted the inherent tendency of fibroblast phenoconversion in HAF culture. TREK-1 expression and activity were higher in fibroblasts compared to myofibroblasts, suggesting a link between TREK-1 and fibroblast phenoconversion.

### 3.3. TREK-1 mRNA expression and fibroblast phenoconversion are interconnected

To test this hypothesis, we investigated the effect of TREK-1 knock-down or overexpression on fibroblast phenoconversion (**Figure 4** a, top). SD-208 and TGF-β1 treatment served as reference points for the fibroblastic and myofibroblastic phenotype, respectively (**Figure 4** a, bottom). Successful TREK-1 knock-down and overexpression were confirmed by qPCR (**Figure 4** b). HAF treated with siTREK-1 expressed more *ACTA2* than siNT treated HAF (1.2 ± 0.2 *vs*. 0.5 ± 0.0) and approached the *ACTA2* mRNA levels of TGF-β1 treated cells (1.5 ± 0.2). HAF overexpressing TREK-1 showed no significant difference in *ACTA2* mRNA expression compared to the empty vector control (eGFP; 1.1 ± 0.1 *vs*. 1.1 ± 0.1; **Figure 4** c). TREK-1 down-regulation also raised *COL1A1* expression compared to siNT treatment (0.8 ± 0.1 *vs*. 0.6 ± 0.0), interestingly TREK-1 overexpression had no significant effect on *COL1A1* expression (0.7 ± 0.0 [TREK-1-eGFP] *vs*. 0.6 ± 0.0 [eGFP]; **Figure 4** d).

**Figure 4.**
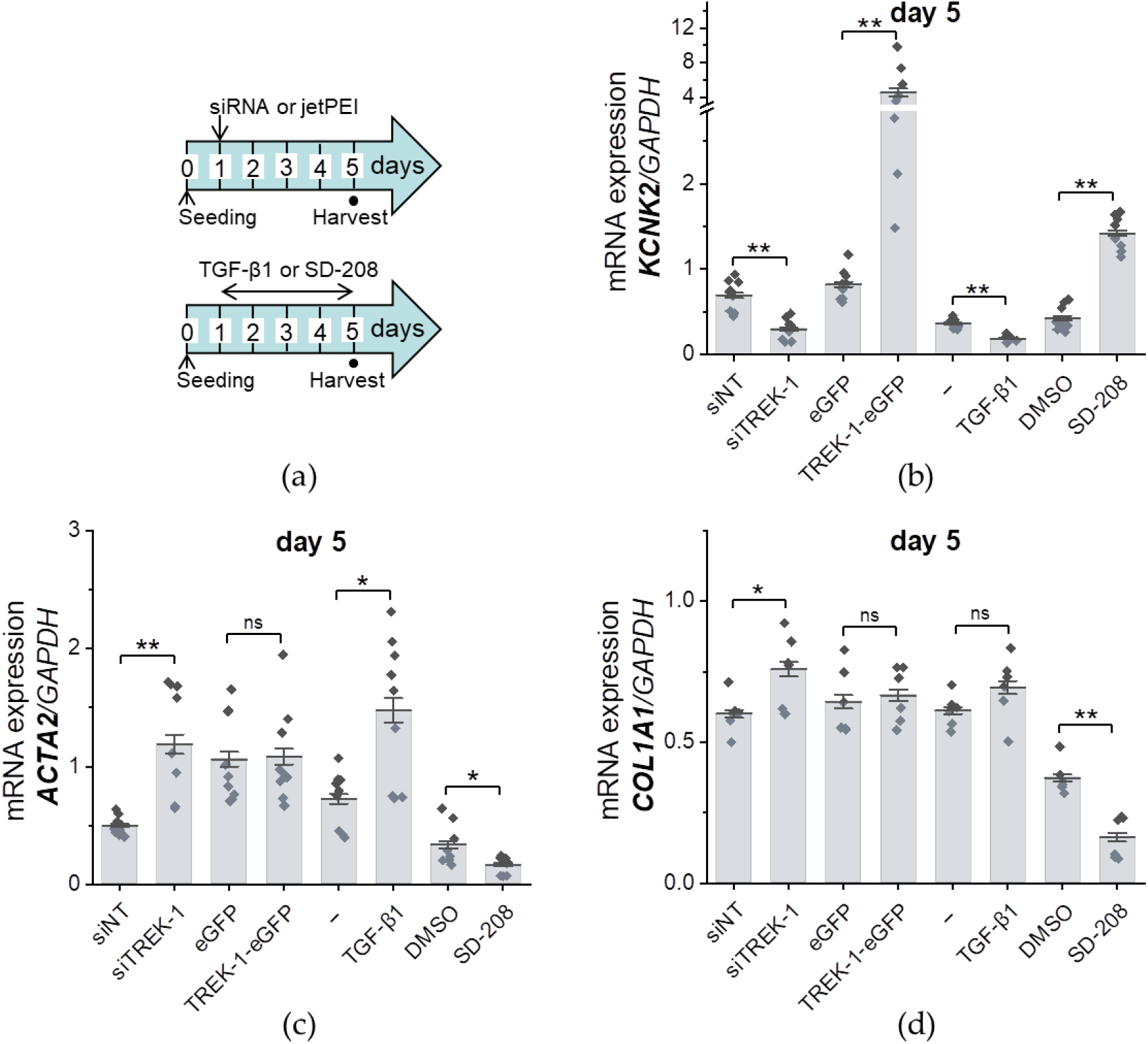
The role of TREK-1 expression in fibroblast phenoconversion; assessed in HAF; qPCR results normalized to *GAPDH*; Mann−Whitney test. (a, top) SiRNA transfection for TREK-1 knockdown (siTREK-1/siT1; non-targeted control: siNT) or jetPEI transfection for TREK-1 overexpression (TREK-1-eGFP; empty vector control: eGFP); (a, bottom) TGF-β1 treatment (400 pmol/L; solvent control: medium only, −) or SD-208 treatment (3 µmol/L; solvent control: 0.03% DMSO). (b−d) *KCNK2* (TREK-1; b), *ACTA2* (αSMA; c) and *COL1A1* (collagen type I, α1 chain; d) mRNA expression on day 5; n = 6−9 dishes/condition; N = 2−3 passages.

In conclusion, *KCNK2* mRNA expression and fibroblast phenoconversion were interconnected. Upon exposure to siTREK-1, HAF express more αSMA and collagen type I, α1 chain than siNT HAF.

### 3.4. TREK-1 expression and activity depend on the heart chamber but not on rhythm status

We then investigated TREK-1 expression and activity in HAFprim. We compared cells outgrown from RA to LA tissue, and samples of patients with SR to those with sustained AF. After 21 days of culture and without passaging, *KCNK2* mRNA expression was higher in HAF_prim_ derived from LA (4.1 ± 0.6) than from RA both in cells from SR and AF patients (4.1 ± 0.6 *vs*. 0.1 ± 0.0, and 2.2 ± 0.5 *vs*. 0.3 ± 0.1 respectively). *KCNK2* mRNA expression was not significantly different between SR and AF from either atrium (**Figure 5** a). To relate changes in *KCNK2* mRNA expression to pro-fibrotic functions, we also assessed *ACTA2* mRNA expression across atria and rhythm backgrounds. *ACTA2* mRNA expression was lower only when comparing RA HAF_prim_ from SR (0.8 ± 0.1) to AF samples (1.2 ± 0.1) (**Figure 5** b).

**Figure 5.**
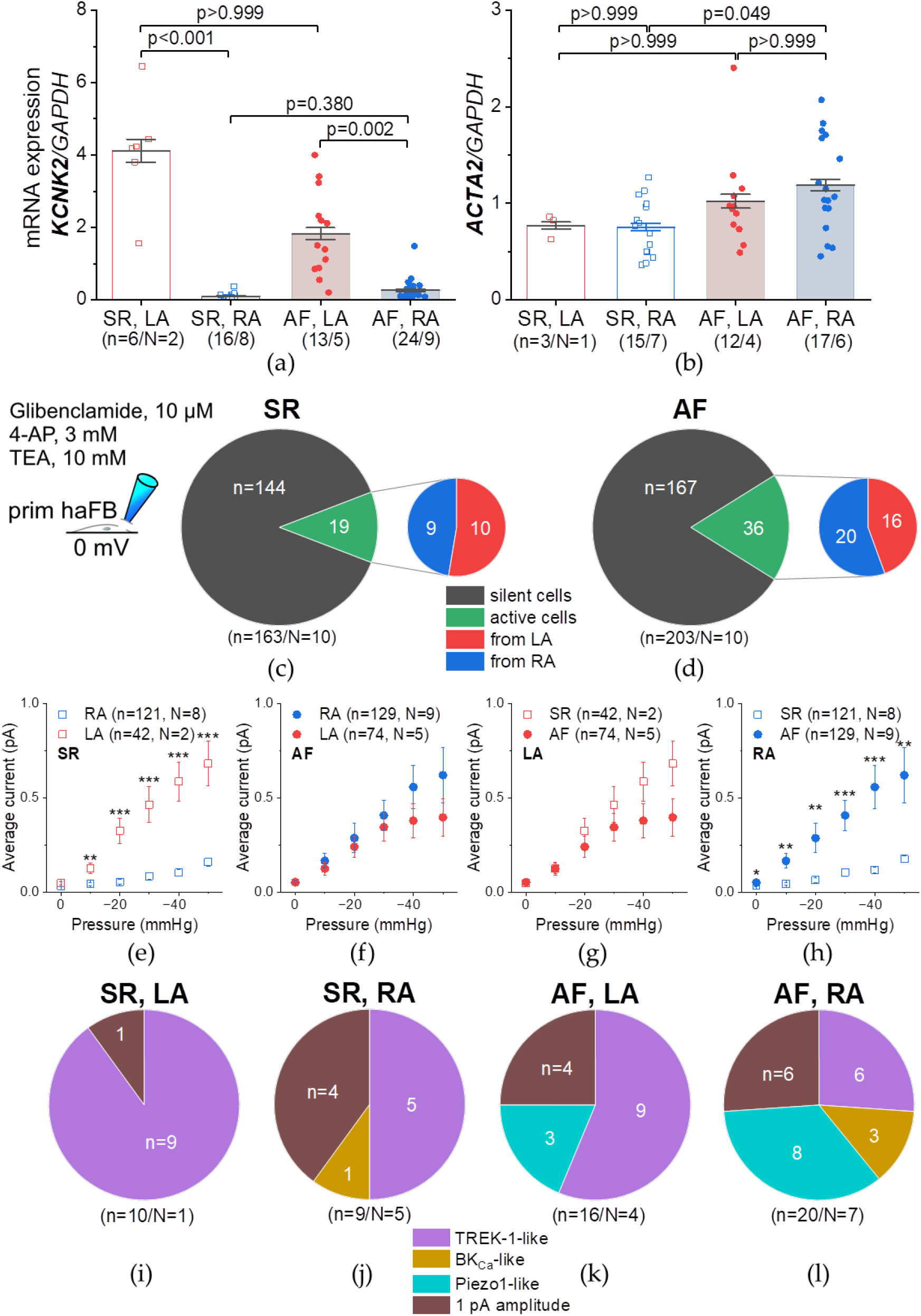
TREK-1 mRNA expression and stretch-induced activity in primary left (LA, red) and right (RA, blue) human atrial fibroblasts (HAFprim) from patients in sinus rhythm (SR, □) or sustained atrial fibrillation (AF, ●). (a, b) qPCR results for *KCNK2* (TREK-1; a) and *ACTA2* (αSMA; b) mRNA expression normalized to *GAPDH*; Kruskal−Wallis ANOVA with Dunn’s *post-hoc* test. (c−h) Patch-clamp measurements in inside-out configuration at a holding potential of 0 mV in the presence of non-K_2P_ channel inhibitors (a, inset). (a, b) Number of silent cells (grey) and active cells (green) in HAF_prim_ derived from patients in SR (c) or AF (d); active cells are further divided into their tissue provenance: LA (red) or RA (blue). (e−h) Average current (over the duration of the pressure pulse)−pressure relationship from 0 to −50 mmHg, comparing RA to LA in SR (e) and AF (f), and comparing SR to AF in LA (g) and RA (h); unpaired t-test. (i−l) Distribution of current patterns; n = number of cells; N = number of patients.

In terms of stretch-induced ion channel activity, the ratios of silent *vs*. active cells are similar when comparing cells from SR and AF patients and LA *vs*. RA (**Figure 5** c, d). Stretch-activated currents were larger in LA than in RA HAF_prim_ from patients in SR, but not in AF (**Figure 5** e, f). RA HAF_prim_ from AF patients show larger currents compared to cells from SR patients (**Figure 5** g, h).

Even though we utilized a patch-clamp protocol and perfusate conditions favoring recordings from TREK-1 channels, we found that other ion channels contributed to current traces. We discriminated four current patterns. The number of TREK-1-like currents was highest in SR LA HAF_prim_ (**Figure 5** i) and lowest in AF RA HAF_prim_ (**Figure 5** l). BK_Ca_- like currents were only observed in RA cells (both SR and AF), whereas Piezo1-like currents were only observed in AF cells (both RA and LA). 1 pA amplitude currents were most frequent in SR RA HAF_prim_ (**Figure 5** j) and lowest in SR LA HAF_prim_ (**Figure 5** i−l).

For four AF patients,, we received RA and LA tissue samples, to compare HAF_prim_ from the same patient. In each individual patient, HAF_prim_ exhibited lower *KCNK2* mRNA expression in RA than LA (**Error! Reference source not found**. a). There was neither a significant difference in *ACTA2* mRNA expression, (**Error! Reference source not found**. b, c), nor in average current at any pressure pulse between RA and LA cells (**Error! Reference source not found**. d−g). The number of active cells, as well as the current amplitudes were inconsistent across patients (**Error! Reference source not found**. d−g). Thus, the paired data confirmed the data presented in **Figure 5**. TREK-1 LA preferential expression was not observed in porcine fibroblasts (paired RA and LA from 4 pigs, **Suppl. Error! Reference source not found**.)

In conclusion, *KCNK2* mRNA expression and activity differ between LA and RA, but do not vary with rhythm status. LA-preferential *KCNK2* mRNA expression was seen in HAF_prim_ derived from patients in SR and AF. At the functional level, LA-preferential TREK-1 activity in was seen only in SR. Neither TREK-1 expression nor activity showed significant differences between SR and AF. The higher average current in RA HAF_prim_ from AF compared to SR samples, was caused by ion channels without characteristic TREK-1 kinetics.

For a broader overview of fibroblast-specific gene expression of ion channels in AF compared to SR samples, we probed the micro-array gene dataset. None of the genes encoding SAC_K_, MSC (**Error! Reference source not found**.), or TREK-1 regulatory partners (**Error! Reference source not found**.) was expressed significantly different between HAF_prim_ from RA of patients in SR or AF (**Error! Reference source not found**. and **Error! Reference source not found**. a).

### 3.5. MSC mRNA expression is chamber- and disease-selective in human hearts

With the aim to identify novel MSC candidates underlying pathological conditions, we conducted a meta-analysis of five previously published bulk RNA-sequencing datasets which included non-diseased cardiac tissue samples and a large number of patients. The mRNA expression of non-diseased and diseased human tissue from ventricles and atria was analyzed with respect to the MSC genes listed in **Error! Reference source not found**..

Cardiac mRNA expression of MSC in non-diseased samples of all heart chambers is shown in **Error! Reference source not found**.. The heatmap represents mRNA expression in normalized counts for each tissue samples and for each MSC gene. It illustrates that MSC have a wide spectrum of mRNA expression in the heart. When we conducted pairwise comparisons, differential gene expression between atria and ventricles accounted for 53% of the variability between individual non-diseased samples, whereas sex-associated differences in gene expression accounted for 10% of variability. In diseased LV samples, principal component 1 roughly separated samples originating from HF patients from those originating from donor hearts (accounting for 22% variability). In RA samples, samples originating from AF patients were intermingled with those originating from SR (**Error! Reference source not found**.).

MSC chamber selectivity was analysed using non-diseased human cardiac tissue samples. We found that 21 MSC displayed chamber-selective gene expression. 17 of these were atria-selective, *i.e*. they are higher expressed in the atria than in the ventricles (**Figure 6** a). For *ASIC1* and *4, CHRNE, KCNA5, KCNMA1, KCNMB2, KCNQ3* and *5, PIEZO2*,

**Figure 6.**
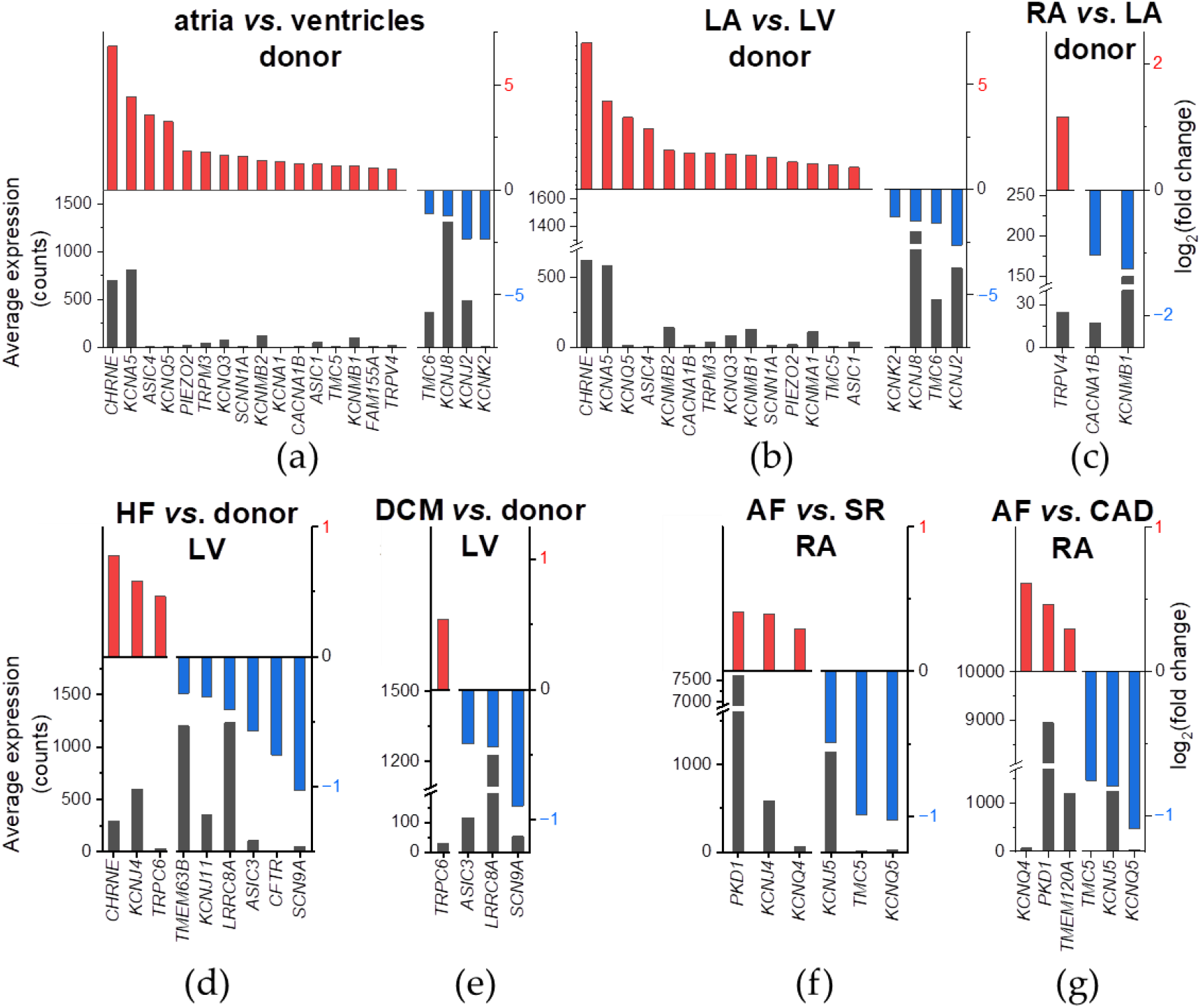
Chamber (a−c)and disease (d−g)-selective MSC mRNA expression in human cardiac tissue samples. Left ordinate indicates the average mRNA expression over all samples from both conditions expressed in normalized counts; right ordinate indicates the log_2_(fold change) of the comparison. MSC genes significantly (adjusted *p*-value < 0.05; Benjamini-Hochberg procedure) higher (left on the abscissa, red) or lower (right on the abscissa, blue) expressed are shown; unit example: log_2_(fold change) = 2 → fold change = 4. (a−c) Atrial (pooled from right [RA] and left [LA] atrium) compared to ventricular (pooled from right [RV] and left [LV] ventricle; a), LA compared to LV (b), and RA compared to LA (c) non-diseased tissue samples. (d, e) Heart failure (HF; pooled from dilated cardiomyopathy [DCM] and ischemic cardiomyopathy; d) and DCM (e) compared to non-diseased LV samples. (f, g) Atrial fibrillation (AF) compared to sinus rhythm (SR; pooled from coronary artery disease [CAD] and heart valve disease; f) and compared to CAD (g) diseased RA samples.

*SCNN1A, TMC5* and *TRPM3* atria selectivity was particularly pronounced in the LA. For *FAM155A* and *KCNA1*, chamber selectivity was particularly strong in the RA. One gene, *KCNMA1*, was expressed higher in LA compared to LV, but showed no significant difference in expression when comparing both atria to both ventricles (**Figure 6** b). *CACNA1B* and *KCNMB1* were clearly identified to be LA-selective. *TRPV4* was clearly identified to be RA-selective (**Figure 6** c). Four MSC were ventricle-selective, *i.e*. they are expressed higher in the ventricles than in the atria (**Figure 6** a). For *KCNK2, KCNJ2* and *8* ventricular selectivity seemed to originate, at least in a major part, from the LV (**Figure 6** b; **Error! Reference source not found**.). At the whole genome level, *KCNA5, CHRNE* and *KCNJ2* ranged among the top 10 differentially expressed genes (**Error! Reference source not found**.).

To address MSC disease selectivity, we compared mRNA expression between diseased and control (non-diseased or SR) human cardiac tissue samples. We found that 15 MSC exhibited disease-selective gene expression. Among those, expression of *CHRNE, KCNJ4* and *TRPC6* was higher and expression of *SCN9A, TMEM63A, KCNJ11, ASIC3, CFTR* and *LRRC8A* was lower in HF samples compared to donor hearts (**Figure 6** d). For *TRPC6, ASIC3, LRRC8A* and *SCN9A*, HF-related differential expression seemed to be driven largely by DCM (**Figure 6** e). In AF, expression of *PKD1, KCNJ4* and *KCNQ4* was higher and expression of *KCNJ5, TMC5* and *KCNQ5* was lower in … compared to samples from patients in SR (**Figure 6** f). Except for *KCNJ4*, this selectivity seemed to be dominated by CAD (**Figure 6** g). One gene, *TMEM120A*, was expressed higher in AF than in CAD, but showed no significant difference in expression when comparing AF to SR (**Figure 6** f, g). At the whole genome level, no MSC ranged among the top 10 differentially expressed genes (**Error! Reference source not found**.).

To conclude this last part of our study, we found that MSC had a wide range of expression in the heart, sex affected gene expression, 21 MSC were expressed chamber selectively and 15 MSC were expressed disease selectively. Interestingly, *KCNQ5* and *CHRNE* were atria-selective and both were downregulated in AF or upregulated in HF, respectively.

## 4. Discussion

In this project we broadened our approach from one ion channel to all known MSC, including MSC candidates, and this from one cell type to cardiac tissue. The overarching aim was to understand better the role of MSC in cardiac pathophysiology.

Our main results obtained in HAF_prim_ are: i) TREK-1 is expressed and active, ii) TREK-1 is more expressed and more active in fibroblasts than in myofibroblasts, iii) TREK-1 knock-down leads to the induction of a myofibroblastic phenotype, iv) TREK-1 shows LA-preferential expression and activity, v) TREK-1 expression and activity are not significantly different in AF compared to SR, and vi) no significant differences in expression of TREK-1 regulatory partners, SAC_K_ and other MSC genes are seen in AF.

Our main results obtained at the whole cardiac tissue level are: i) TREK-1 expression is ventricle-selective and not significantly different between non-diseased and HF, or between SR and AF, ii) expression of acetylcholine receptor nicotinic subunit ε is upregulated in atria (compared to ventricles) 21 of 54 and in HF (compared to donor), iii) TRPV4 is expressed RA selectively, iv) Cav2.2 and BKCa β-1 subunit are expressed LA selectively, v) Kir2.3 is upregulated and Nav1.7 is downregulated in HF, and vi) TRPP1 is upregulated and GIRK4 is downregulated in AF.

### 4.1. TREK-1 is present but lowly expressed in HAFprim, and implicated in fibroblast phenoconversion

We showed TREK-1 mRNA expression and activity in HAF_prim_ (**Figure 1** and **Figure 2** for HAF; **Figure 5** and **Error! Reference source not found**. for HAFprim). With this, our study is the first report on TREK-1 activity in HAFprim. In comparison to other AA-sensitive ion channels and MSC, TREK-1 mRNA expression was low. For example, BK_Ca_ was also expressed and active in HAF_prim_ (**Error! Reference source not found**. and **Error! Reference source not found**.), as reported before [63]. Only a small number of HAF_prim_ displayed TREK-1 activity. In HAF, TREK-1 activity was small and inter-cell variability was large because the cell line contained a mixed population of fibroblasts and myofibroblasts, with a majority of myofibroblasts. Indeed, we showed higher expression and activity of TREK-1 in fibroblasts compared to myofibroblasts (**Figure 2**). The difference in TREK-1 mRNA expression between fibroblasts and myofibroblasts was reversible in both directions (**Figure 3**). Similarly, lower percentage of Na_v_1.5 positive cells in HAF_prim_ [92] and lower SUR2 and K_ir_6.1 protein expression in mouse ventricular fibroblasts [93] compared to myofibroblasts has been reported before.

We investigated whether TREK-1 could influence fibroblast phenoconversion. We observed higher αSMA and collagen type I α1 chain mRNA expression levels in siTREK-1 than in siNT treated HAF (**Figure 4**). This suggests that TREK-1 has anti-fibrotic properties and thus, plays a role in fibroblast phenoconversion and fibrotic tissue remodelling. However, this finding does not match an earlier report on reduced fibrosis in mechanically overloaded mice hearts with fibroblast-specific TREK-1 knock-out [17]. It would be interesting to identify whether the difference in species (murine *vs*. human) or in chambers (ventricles *vs*. atria) could explain the apparent discrepancy. Overexpression of TREK-1 did not significantly influence *ACTA2* or *COL1A1* mRNA expression (**Figure 4**), possibly due to the low (≈ 5%) transfection success rate, or to a one-way mechanism where only TREK-1 downregulation affects the mRNA expression of myofibroblastic marker genes.

### 4.2. Modulation of fibroblast phenoconversion

Our results highlight that it is crucial to assess and control fibroblast phenoconversion in cell culture conditions and thereby define the studied population of cells. With SD-208 and TGF-β1 we were able to efficiently modulate the fibroblast / myofibroblast balance in either direction irrespective of the concomitant inherent trends of phenoconversion in HAF culture. As HAF proliferated between day 1 and day 7 of culture, there was a general tendency towards a more fibroblastic phenotype. Between day 7 and day 13, HAF became overcrowded which may have led to cell death, consequently lower cell density and enhanced differentiation into myofibroblasts. In fact, differences in cell density can influence fibroblast phenoconversion [94], *e.g*. culture of murine ventricular fibroblasts at high density resulted in fibroblasts, whereas they differentiated myofibroblasts when seeded at low density [95]. In line with our observations, others reported that HAF_prim_ and mouse ventricular fibroblasts cultured for 7 days were fibroblastic and they differentiated into myofibroblasts between 7 and 12 days of culture [92, 93].

SD-208 solvent control exposure (0.03% DMSO) caused independent effects on *KCNMA1* mRNA expression. It has been reported before that DMSO can induce changes in gene expression [96]. Thus, a potential effect of DMSO itself must be considered. Moreover, *COL3A1* mRNA expression was higher in HAF treated with SD-208 compared to DMSO. Indeed, *COL3A1* mRNA expression has been shown to be either positively [97, 98] or negatively [99–102] correlated with the presence of myofibroblasts.

During the experiments to assess phenotype reversibility, we observed that ≈85% of TGF-β1 pre-treated HAF detach upon SD-208 treatment between day 7 and day 13. Since, no cell detachment was observed between day 1 and day 7, this may have been due to cell overcrowding. In addition, we do not know whether the phenotype of one individual cell was reversible or whether we modulated differential proliferation of a distinct phenotype. Taken together, the possibility for reversibility remains to be further clarified.

### 4.3. TREK-1 modulation by heart chamber and rhythm status

In HAF_prim_ derived from patients in SR, we showed LA-preferential TREK-1 expression and activity. In HAF_prim_ from patients with AF, this difference in expression remained, but no significant differences in stretch-induced functional activity were seen. This may be explained by a reduced contribution of TREK-1 to the overall current in AF LA HAF_prim_ (**Figure 5**). Higher LA than RA whole tissue TREK-1 mRNA expression in mice but not in humans was reported before [25]. Here, we show this LA-preferential human TREK-1 mRNA expression to be fibroblast-specifically. The overall low number in active HAF_prim_ reflects the ventricle/LV-selective TREK-1 mRNA expression observed at the whole tissue level (**Figure 6**). This confirms an earlier report on higher human TREK-1 mRNA expression in LV compared to RA and to LA [25].

Our results did not show a significant differential regulation of TREK-1 mRNA expression or activity associated with AF. The higher average current in RA HAF_prim_ of AF patients compared to SR patients did not display the characteristic TREK-1 kinetics (**Figure 5**). We believe that other ion channels contaminated our recordings. In fact, Piezo1 activity has been shown to be upregulated in HAF_prim_ from patients in AF [63]. Previously, TREK-1 mRNA downregulation was reported in fresh LV, RA (mRNA and protein) and LA (mRNA) tissue of patients in AF (with or without concomitant HF) [25, 26]. A possible explanation for this discrepancy may be that the AF phenotype in our HAF_prim_ had vanished by the time of outgrowth and by other artefacts of cell culture. For instance, in [63], we show that Piezo1 mRNA expression decreases over passages. For future experiments, freshly isolated HAF_prim_ will allow capturing a close-to-*in vivo* status.

Expression of αSMA was lower in RA HAF_prim_ from SR than from AF patients but did not significantly differ between RA and LA (**Figure 5**). Characterization of tissue derived from patients will be expanded by quantification of tissue stiffness and of fibrosis.

In this study, patients with AF were significantly older than patients in SR. Thus, we do not only compare between rhythm status but also between age groups which is a major limitation of this study. Furthermore, we only discriminate between RA and LA but we cannot specify whether tissue samples were obtained from the appendage or the free wall. However, ion channel presence may differ between these locations.

### 4.4. MSC mRNA expression in human cardiac health and disease

The chamber-selective TREK-1 mRNA expression and activity as well as the role of TREK-1 in fibroblast phenoconversion were intriguing with regard to the approach of chamber-selective drug therapy. In the most general approach, we sought to characterise MSC mRNA expression in human cardiac health and disease with the aim to identify chamber-selective drug targets.

From heatmap data, we conclude that most MSC are widely expressed within the heart (**Error! Reference source not found**.). That said, we should not forget that minimally expressed MSC may still play a major role in cardiac physiology [103]. Comparing the expression of individual MSC in non-diseased atrial tissue to their expression in HAF_prim_ from SR patients, we observed differences in the abundance of distinct MSC. Thus, *TMC3, KCNQ5* and *KCNK4* were lowly expressed in atria, but displayed high fibroblast-specific expression. The opposite holds true for *KCNJ8, KCNQ1* and *CACNA1C* (**Error! Reference source not found**.).

PCA showed that differential gene expression is determined by heart chamber, sex, and health status. The samples originating from AF from those originating from SR may be explained by the fact that all patients with AF also had CAD or HVD as reason for cardiac surgery (just as SR patients) but tissue samples were only grouped by atrial rhythm. In addition, all diseased RA samples were obtained from male patients. Thus, no sex-associated differences could be explored (**Error! Reference source not found**.).

Differentially expressed genes in non-diseased human cardiac tissue derived from different heart chambers are summarized in **Error! Reference source not found**.. Comparing LA to LV, we confirmed *KCNA5* upregulation and *KCNJ2* downregulation but we could not confirm *SCN5A, SCN9A* and *KCNJ5* upregulation in LA when compared to LV [104, 105].

Differentially expressed genes in cardiac tissue derived from patients with different health conditions are summarized in **Error! Reference source not found**.. We confirmed *CFTR* [106, 107] and *KCNJ11* [108] downregulation in HF as well as *TRPC6* [12] upregulation in DCM. Interestingly, among the MSC differentially expressed between AF and SR at the whole tissue level (**Figure 6**), none recurred in HAF_prim_ (**Error! Reference source not found**.). In general, we found that MSC are lowly regulated in cardiac disease. However, as for ion channels, it may not be a change in expression but a change in activity that contributes to cardiac pathophysiology. For example, in atrial myocytes, K_v_1.5 has been shown to respond to shear stress by increased insertion into the plasma membrane, *i.e*. recruitment from intracellular compartments [109].

The top 10 differentially expressed genes associated to heart chamber or cardiac diseases confirm earlier findings which contribute to validate our study (**Error! Reference source not found**. and **Error! Reference source not found**.). Differentially expressed genes below the set threshold may still contribute to the pathophysiology of a certain disease.

Whole tissue RNA-sequencing does not allow for interpretation about the cell type responsible for differential gene expression and for quantification of heterogeneous responses in individual cell populations. Most functional data were acquired in a specific cell type (often cardiomyocytes). The obtained results will be advanced by single-cell or single-nucleus RNA-sequencing. Moreover, qPCR and functional experiments like patch-clamp can be used to confirm expression and characterise candidate MSC genes. Furthermore, the genetic background was not consistent within the dataset but could not be accounted for in the analysis, in part, because the information could not be retrieved from all original studies.

This meta-analysis gives a comprehensive overview of MSC mRNA expression in cardiac health and disease, and highlights MSC chamber preferential expression. By identifying candidate genes which show chamber- and/or disease selectivity at the mRNA level, novel potential molecular targets involved in mechano-transduction are featured.

## 5. Conclusions

MSC represent a class of mechanoreceptors differentially expressed in various cardiac diseases. This report highlights the broad expression of MSC in the heart in health and diseases, in general, and the implication of TREK-1 in phenoconversion with potential relevance for collagen production, in particular.

Based on the data, the key conclusions are:

i. TREK-1 is expressed and active in HAFprim. Its activity and expression depend on the heart chamber but not on the rhythm status.
ii. TREK-1 is higher expressed and active in fibroblasts than in myofibroblasts. Its down regulation leads to a myofibroblastic phenotype suggesting a role in phenoconversion.
iii. *CACNA1B* and *KCNMB1* are expressed LA selectively (compared to LA); TRPV4 is expressed RA selectively (compared to LA); CHRNE is atria-selective (compared to ventricles) and upregulated in HF tissue (compared to donor tissue); KCNJ4 is up- and SCN9A is downregulated in HF tissue (compared to donor tissue); PKD1 is up- and KCNJ5 is downregulated in AF tissue (compared to SR tissue)..

This work identifies potential mechanisms and actors of pathological tissue remodelling in the course of different cardiac diseases, where fibrosis is often associated to impaired contractile function.

## Supplementary Materials

The following are available online at www.mdpi.com/xxx/s1.

- Figure S1: Proposed classification of MSC.
- Figure S2: AA-induced currents in HAF.
- Figure S3: AA- and spadin-induced currents in HAF.
- Figure S4: Expression and activity of AA-sensitive ion channels in HAFprim.
- Figure S5: Paired mechanically-induced TREK-1 activity of transfected HEK cells in cell-attached and inside-out configuration.
- Figure S6: Single channel amplitude and E_rev_ of TREK-1 in transfected HEK cells.
- Figure S7: Cell area and percentage of αSMA positive HAF differentiated into (myo)fibroblasts.
- Figure S8: mRNA expression of genes linked to fibrosis in HAF differentiated into (myo)fibroblasts.
- Figure S9: mRNA expression of genes encoding SAC in HAF differentiated into (myo)fibroblasts.
- Figure S10: Paired TREK-1 expression and stretch-induced activity in RA and LA HAF_prim_ from patients in AF.
- Figure S11: Paired TREK-1 expression and stretch-induced activity in primary RA and LA porcine atrial fibroblasts.
- Figure S12: Gene expression of SAC_K_ and TREK-1 regulatory partners in HAF_prim_ obtained from patients in SR or AF.
- Figure S13: MSC mRNA expression in HAF_prim_ obtained from patients in SR, or in non-diseased human cardiac tissue samples.
- Figure S14: Principal component analysis.
- Figure S15: Chamber-selective mRNA expression in non-diseased human cardiac tissue samples.
- Figure S14: Disease-selective mRNA expression in LV and RA human cardiac tissue samples.
- Table S1: Human MSC.
- Table S2: TREK-1 regulatory partners.
- Table S3 and S4: Summary of chamber-selective MSC gene expression in non-diseased human cardiac tissue samples.
- Table S5: Summary of disease-selective differential MSC expression.

## Author Contributions

Conceptualization, ED, UR and RP; methodology, ED, DY, RB and RP; formal analysis, ED and DY; investigation, ED; data curation, ED; writing—original draft preparation, ED; writing—review and editing, ED, PK, UR and RP; visualization, ED; supervision, UR and RP; project administration, RP; funding acquisition, PK, UR and RP. All authors have read and agreed to the published version of the manuscript.

## Institutional Review Board Statement

The study was conducted according to the guidelines of the Declaration of Helsinki, and approved by the Ethics Committee of the University of Freiburg, Freiburg im Breisgau, Germany (CardioVascular BioBank ethical approval: 393/16; study approval: 10719), the Medical Faculty of the Technical University Dresden, Dresden, Germany (ethical approval: 15.1/01031/006/2008 and EK790799), the Medical Faculty of Heidelberg University, Heidelberg, Germany (ethical approval: S-017/2013) and the Medical Research Council Budapest, Scientific and Research Ethics Committee, Budapest, Hungary (study approval: 4991-0/2010-1018EKU [339/PI/010]).

## Informed Consent Statement

Informed consent was obtained from all subjects involved in the study.

## Data Availability Statement

The datasets presented in this study can be found in online repositories. The names of the repositories and accession numbers can be found below:

- https://www.ncbi.nlm.nih.gov/geo/query/acc.cgi?acc=GSE57345
- https://www.ncbi.nlm.nih.gov/geo/query/acc.cgi?acc=GSE71613
- https://www.ncbi.nlm.nih.gov/geo/query/acc.cgi?acc=GSE112339
- https://www.ncbi.nlm.nih.gov/geo/query/acc.cgi?acc=GSE128188
- https://www.ebi.ac.uk/ena/browser/view/PRJEB42485

## Acknowledgments

We thank the staff, especially Fabian A. Kari, and patients of the Department of Cardiovascular Surgery of the University Heart Center Freiburg·Bad Krozingen for access to human tissue. We are grateful to the staff of CardioVascular BioBank in Freiburg im Breisgau for providing human tissue samples. We thank Constanze Schmidt and Felix Wiedmann from the Department of Cardiology, University Hospital Heidelberg, Heidelberg, Germany for providing ventricular tissue samples. We acknowledge the support of the Medical Faculty Carl Gustav Carus, TU Dresden, Germany for providing some atrial tissue samples. We are grateful to István Baczkó from Department of Pharmacology and Pharmacotherapy, University of Szeged, Szeged, Hungary for providing non-diseased tissue samples. We thank Eric Honoré and Delphine Bichet for the gift of the TREK-1-IRES2-eGFP plasmid, and Frederick Sachs for the Piezo1-IRES2-eGFP plasmid. We acknowledge the support of the Freiburg Galaxy Team: Dr. Björn Grüning, Bioinformatics, University of Freiburg (Germany) funded by the Collaborative Research Centre 992 Medical Epigenetics (DFG grant SFB 992/1 2012) and the German Federal Ministry of Education and Research (BMBF) grant 031 A538A de.NBI-RBC. We also thank Tony Matschulla from the Institute of Experimental and Clinical Pharmacology and Toxicology, Faculty of Medicine, University of Freiburg, and Eike M. Wülfers from the Institute for Experimental Cardiovascular Medicine, University Heart Center Freiburg-Bad Krozingen for their technical support. ED, DY, RB, PK, UR, and RP are members of SFB1425, funded by the German Research Foundation (DFG, Project #422681845). UR is member of the steering committee of the Atrial Fibrillation NETwork (AFNET), Münster, Germany.

