## Supplemental figures for "TREK-1 knock-down in human atrial fibroblasts leads to a myofibroblastic phenotype: a role in phenoconversion and overview of mechano-sensitive channel mRNA expression in cardiac diseases"

*Article*

### SUPPLEMENTARY FIGURES

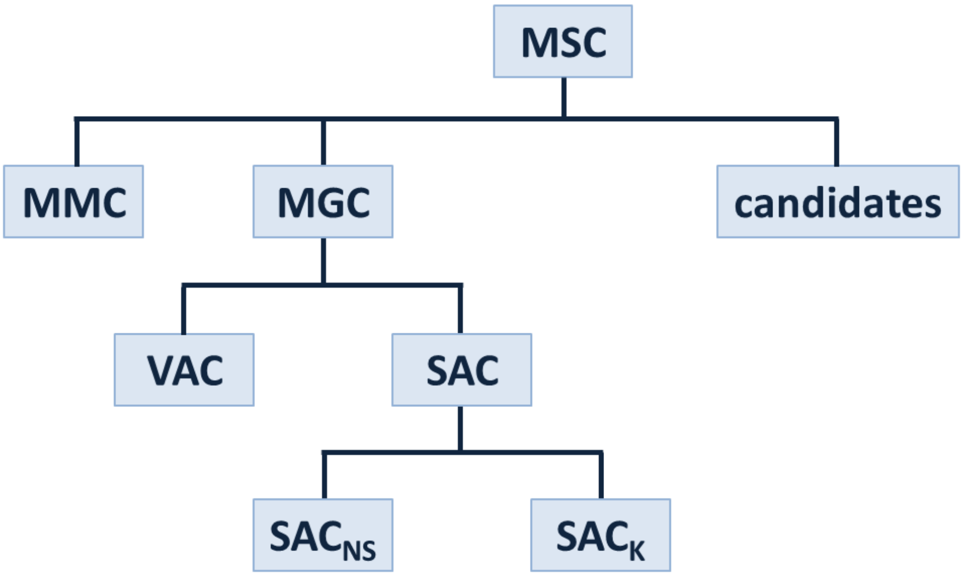


**Suppl. Figure 1.** Proposed classification of mechano-sensitive ion channels (MSC), including mechanically modulated (MMC) and mechanically gated ion channels (MGC). MGC include cell volume-activated ion channels (VAC), and stretch-activated ion channels (SAC). The latter are subdivided into cation non-selective $SAC_{NS}$ and $K^+$-selective $SAC_K$. Additionally, there are MSC candidates, referring to ion conducting pathways that have not yet been shown to be mechano-sensitive, but which – based on phylogenetic analysis [110] and sequence similarity [111], *e.g.* number of transmembrane domains of the corresponding proteins and/or links with mechano-transduction pathways – are predicted to be mechano-sensitive.

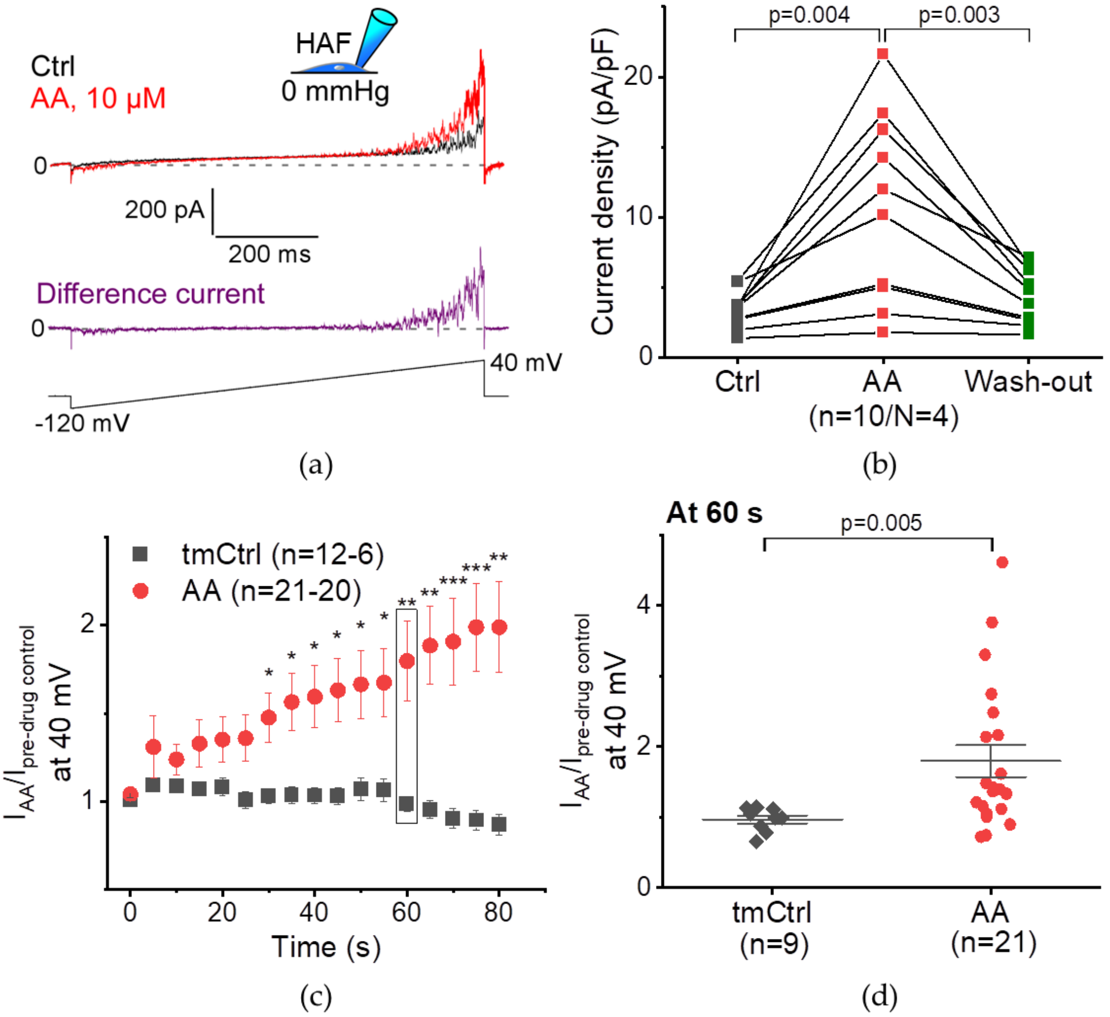


**Suppl. Figure 2.** Arachidonic acid (AA, red)-induced currents in immortalized human atrial fibroblasts (HAF). Patch-clamp measurements in whole-cell configuration (pipette pressure 0 mmHg; a, inset) in response to a 800 ms long ramp protocol from −120 to 40 mV (a, lowest trace). (a) Representative recording of absolute current; top trace: pre-drug control (Ctrl, black) and after 80 s of exposure to AA (10 µmol/L); middle trace: AA-sensitive current measured as difference current (AA – Ctrl, purple). (b) Current density amplitudes (expressed in pA/pF) at 40 mV before (Ctrl), after exposure to AA, and after wash-out of AA (green); Wilcoxon signed rank test. (c) Time course of current relative to Ctrl value ($I_{AA}/I_{pre\text{-}drug\ control}$) at 40 mV for cells exposed to AA or to bath solution only (time-matched control; tmCtrl, black); n-numbers decrease during the protocol. (d) Single data points at 60 s of exposure from the graph in c; comparison of AA-induced current to tmCtrl; (c, d) Mann−Whitney test.

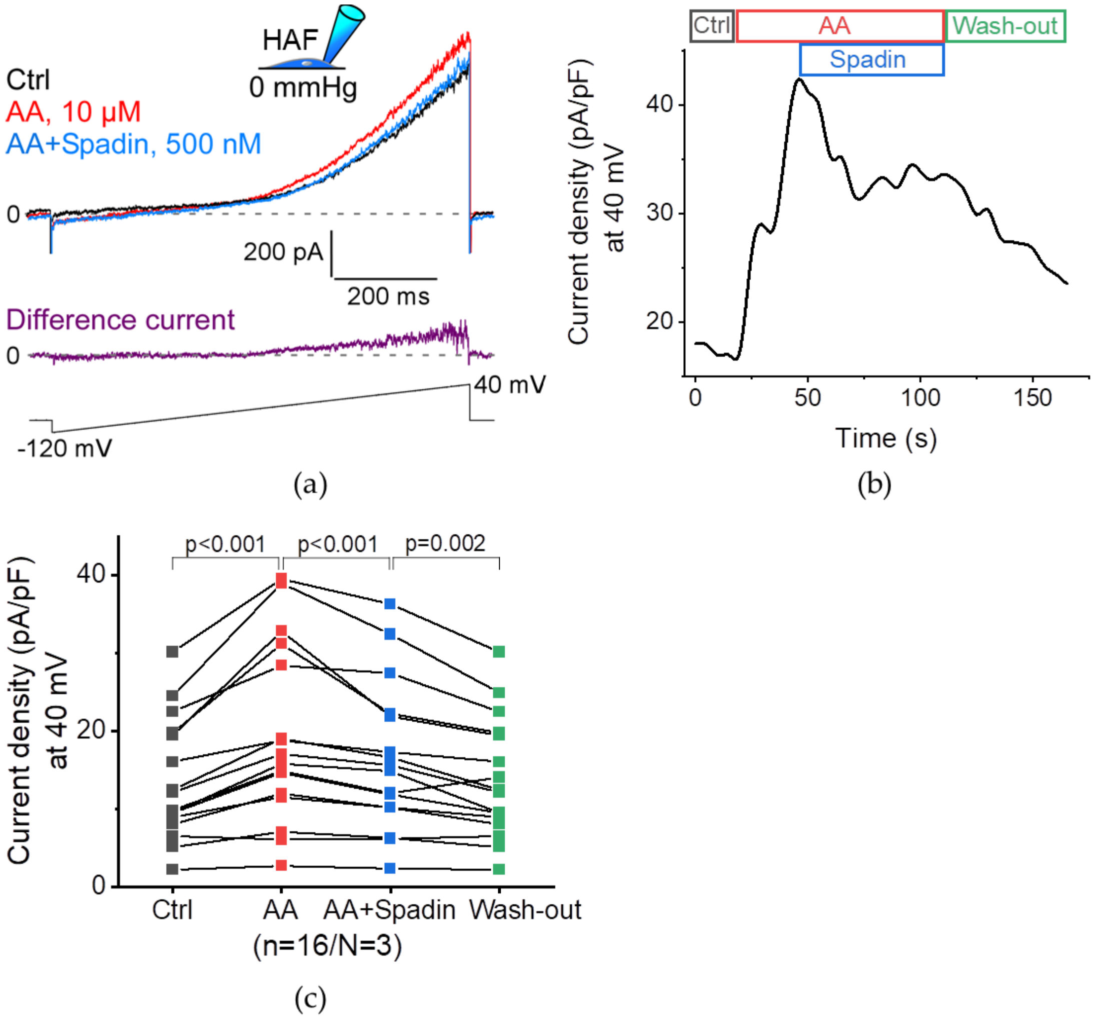


**Suppl. Figure 3.** Arachidonic acid (AA, red)- and spadin (blue)-induced currents in immortalized human atrial fibroblasts (HAF). Patch-clamp measurements in whole-cell configuration (pipette pressure 0 mmHg; a, inset) in response to a 800 ms long ramp protocol from −120 to 40 mV (a, lowest trace). (a) Representative recording of absolute current; top trace: pre-drug control (Ctrl, black), in the presence of AA (10 µmol/L) and in the presence of AA and spadin (500 nmol/L); middle trace: Spadin-sensitive current measured as difference current (AA+Spadin – Ctrl, purple). (b) Representative recording of current density (expressed in pA/pF) at 40 mV over time; top: perfusion protocol with Ctrl, AA, spadin and wash-out (green). (c) Current density amplitudes at 40 mV before (Ctrl), after exposure to AA, after exposure to AA and spadin, and after wash-out of AA; Paired sample Wilcoxon signed rank test. Results: AA increased current amplitude from 13.5 ± 1.9 pA/pF to 19.4 ± 2.9 pA/pF. This effect was reversed upon additional exposure of spadin (to 16.4 ± 2.4 pA/pF). The current density was further decreased by wash-out of AA and spadin (5.9 ± 1.1 pA/pF).

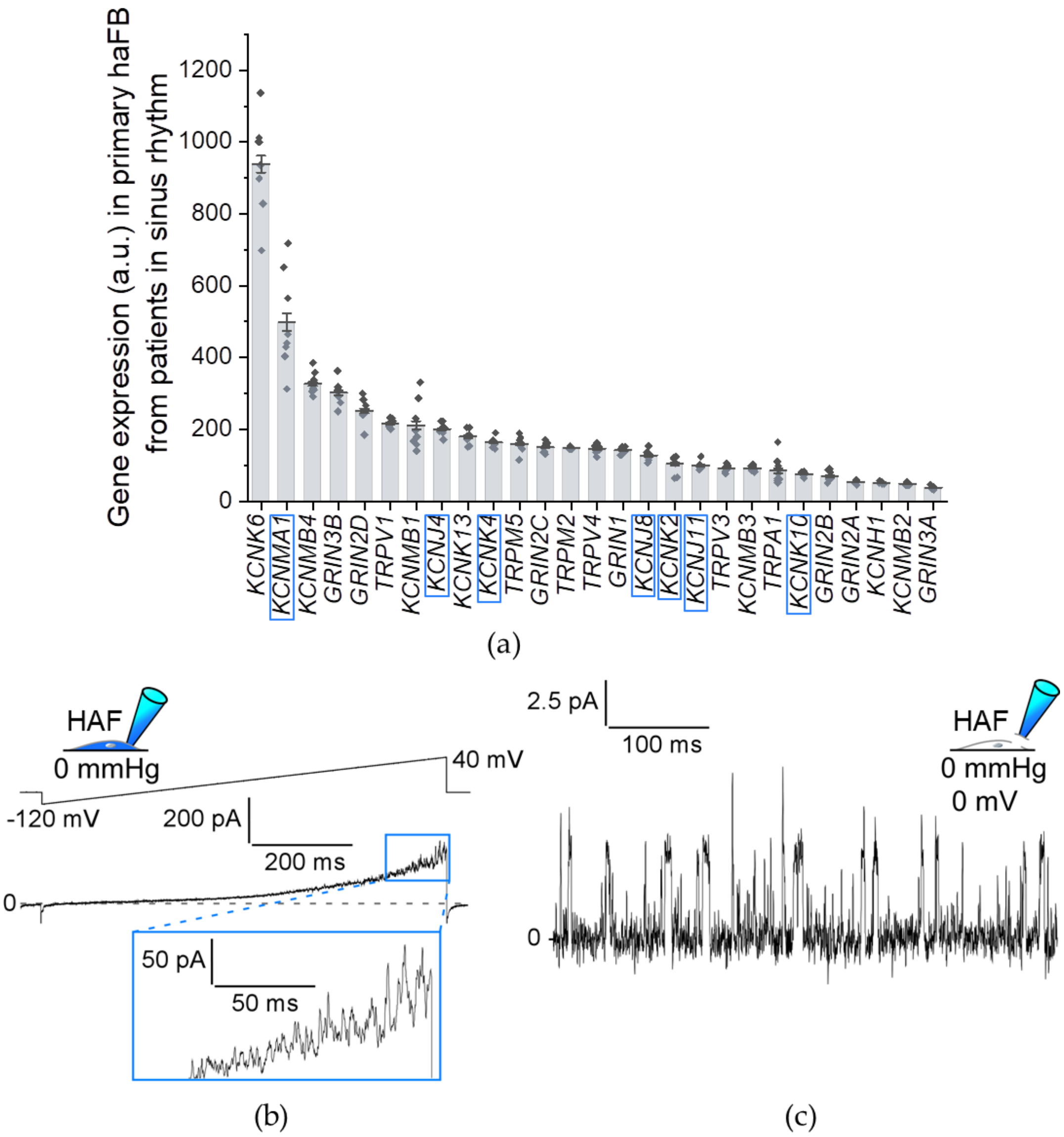


**Suppl. Figure 4.** Expression and activity of arachidonic acid (AA)-sensitive ion channels in human atrial fibroblasts (HAFprim). (a) Gene expression in $HAF_{prim}$ from 8 patients with sinus rhythm; assessed by gene micro-array; arbitrary unit (a.u.). AA-sensitive ion channels are $K_{ir}2.3$ [112], the voltage-gated $K^+$ channel EAG1 [113], the inward rectifier $K_{ATP}$ channels $K_{ir}6.1$ and 6.2 [114], $BK_{Ca}$ [115], the $K_{2P}$ channels TRAAK [64], TWIK-2 [116], TREK-1 and TREK-2 [15], the small-conductance background $K^+$ channel THIK-1 [117], TRPV1 [118], TRPV3 [119], TRPV4 [120], ankyrin TRP (TRPA)1 [121], TRPM2 [122], TRPM5 [123], and ionotropic glutamate receptor NMDA [124]. Genes for $K^+$-selective stretch-activated ion channels are highlighted in blue. They were expressed at the following levels: $BK_{Ca}$ subunit $\alpha$-1 (*KCNMA1*, 498.0 ± 48.0) [125], $K_{ATP}$ (*KCNJ8*, 126.9 ± 5.0; *KCNJ11*, 99.5 ± 3.9), inward rectifier $K^+$ channel 4 (*KCNJ4*) and the $K_{2P}$ channels TREK-1 (*KCNK2*, 103.9 ± 8.7), TREK-2 (*KCNK10*, 75.2 ± 2.0) and TRAAK (*KCNK4*, 163.2 ± 4.7). (b, c) $BK_{Ca}$ channel-like currents in immortalized $HAF_{prim}$ (HAF), determined by patch-clamp measurements. (b) Representative recording of outward current in whole-cell configuration (pipette pressure 0 mmHg; a, inset) in response to a ramp protocol from −120 to 40 mV in 800 ms (a, top trace). (c) Representative recording of single channel activity in inside-out configuration (pipette pressure 0 mmHg; holding potential 0 mV; b, inset).

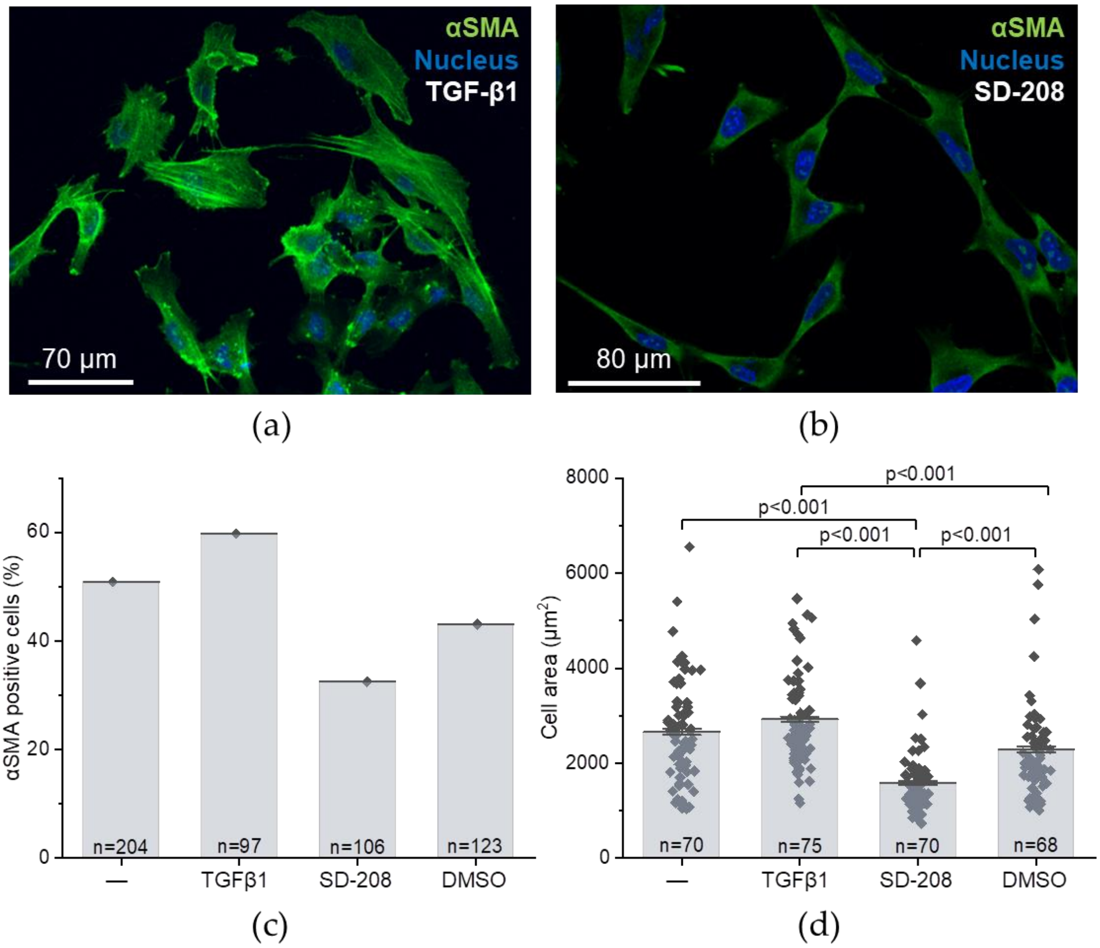


**Suppl. Figure 5.** Cell area and percentage of αSMA positive HAF, determined by confocal microscopy. Cell culture medium only (−), or supplemented with TGF-β1 (400 pmol/L) or SD-208 (3 µmol/L; solvent control: 0.03% DMSO). (a, b) Representative images with αSMA in green and nuclei in blue. (c) Quantification of αSMA-positive cells based on the fluorescence intensity; N = 1 passage; in decreasing order: 59.8% for TGF-β, 50.9% for medium only, 43.1% for DMSO, 32.5% for SD-208. (d) Single data points for cell area; one data point represents one cell; N = 1−2 passages; Kruskal−Wallis ANOVA with Dunn's *post-hoc* test (only significant differences are indicated).

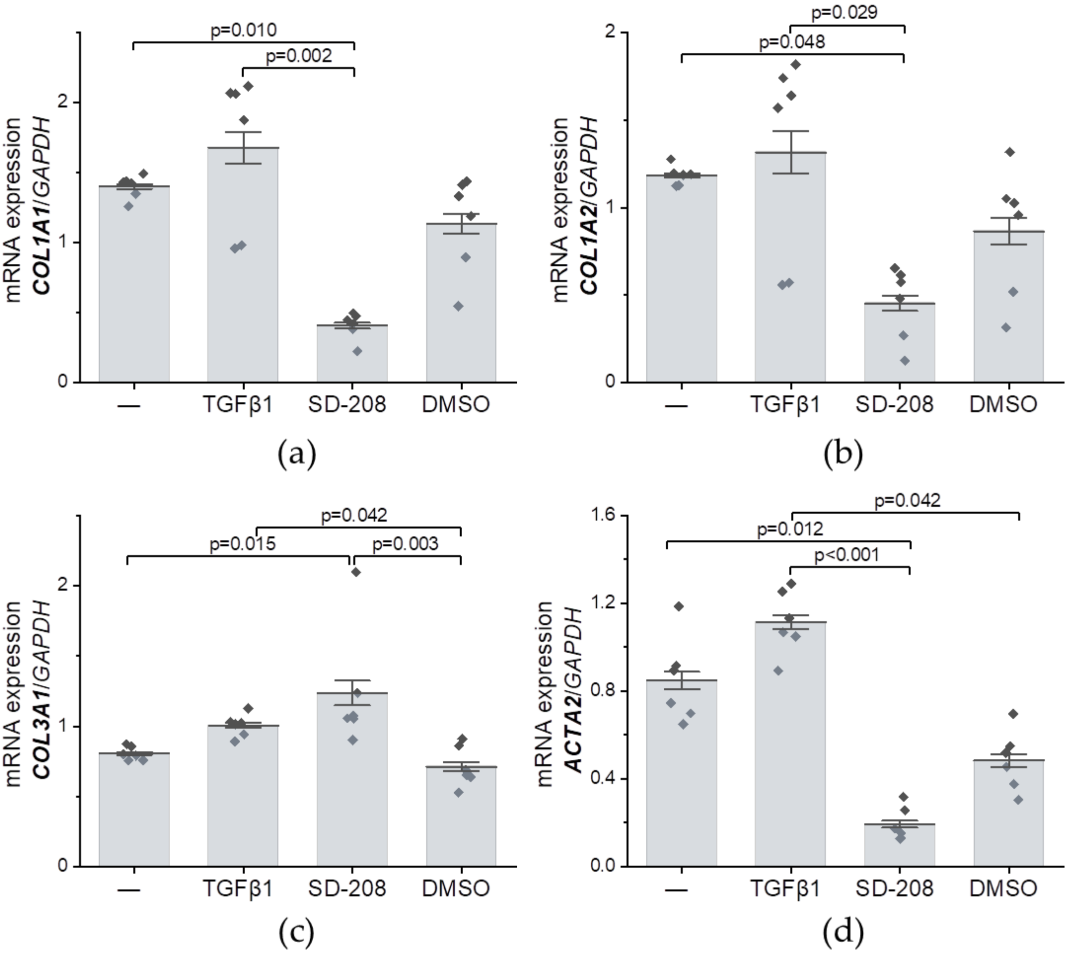


**Figure 6.** mRNA expression of genes linked to fibrosis in HAF, determined by qPCR and normalized to the expression of *GAPDH* (housekeeping gene). Cell culture medium only (−), or supplemented with TGF-β1 (400 pmol/L) or SD-208 (3 µmol/L; solvent control: 0.03% DMSO) were compared; n = 6 wells, N = 3 passages for all conditions; Kruskal−Wallis ANOVA with Dunn's *post-hoc* test (only significant differences are indicated). (a) gene: *COL1A1* (protein: collagen type I, α1 chain). (b) *COL1A2* (collagen type I, α2 chain). (c) *COL3A1* (collagen type III, α1 chain). (d) *ACTA2* (smooth muscle actin).

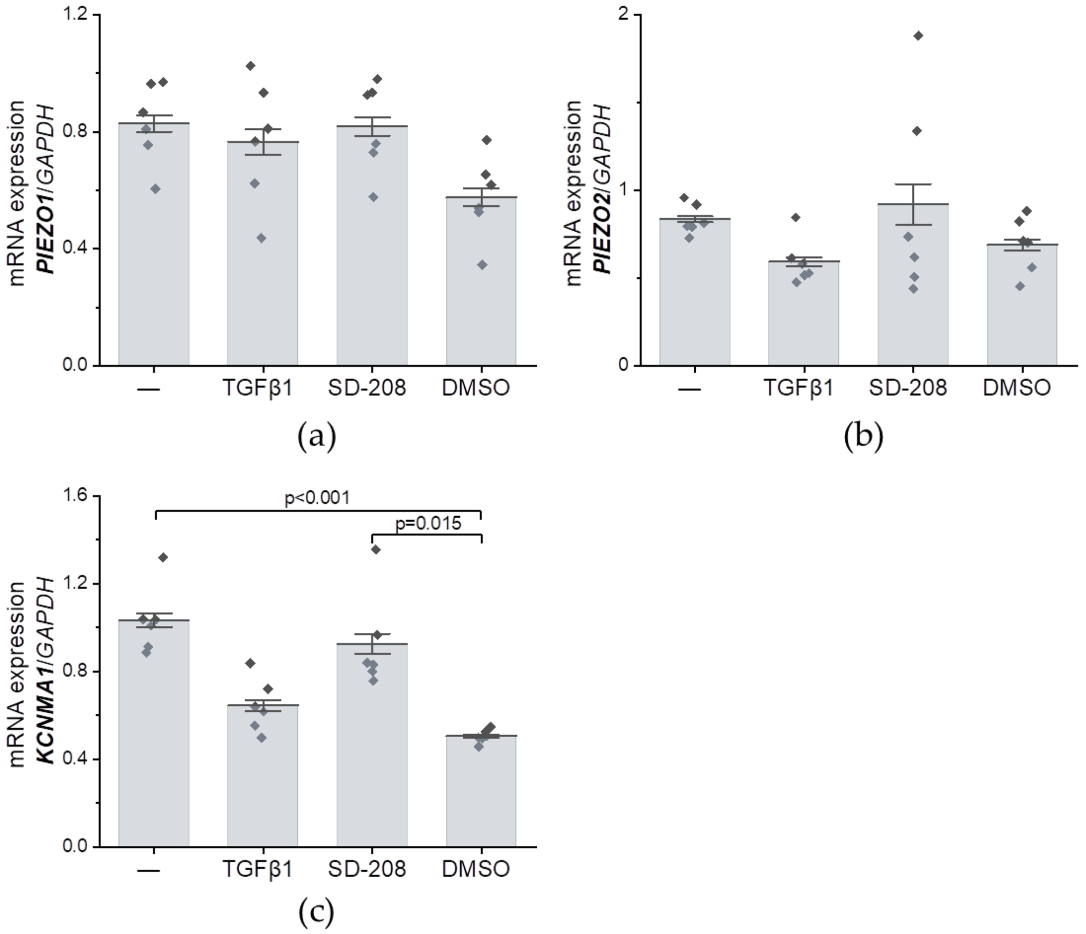


**Suppl. Figure 7.** mRNA expression of genes encoding SAC in HAF, determined by qPCR and normalized to the expression of *GAPDH* (housekeeping gene). Cell culture medium only (−), or supplemented with TGF-β1 (400 pmol/L) or SD-208 (3 µmol/L; solvent control: 0.03% DMSO) were compared; n = 6 wells, N = 3 passages for all conditions; Kruskal−Wallis ANOVA with Dunn's *post-hoc* test (only significant differences are indicated). (a) *PIEZO1* (Piezo1). (b) *PIEZO2* (Piezo2). (c) *KCNMA1* ($BK_{Ca}$ $\alpha$).

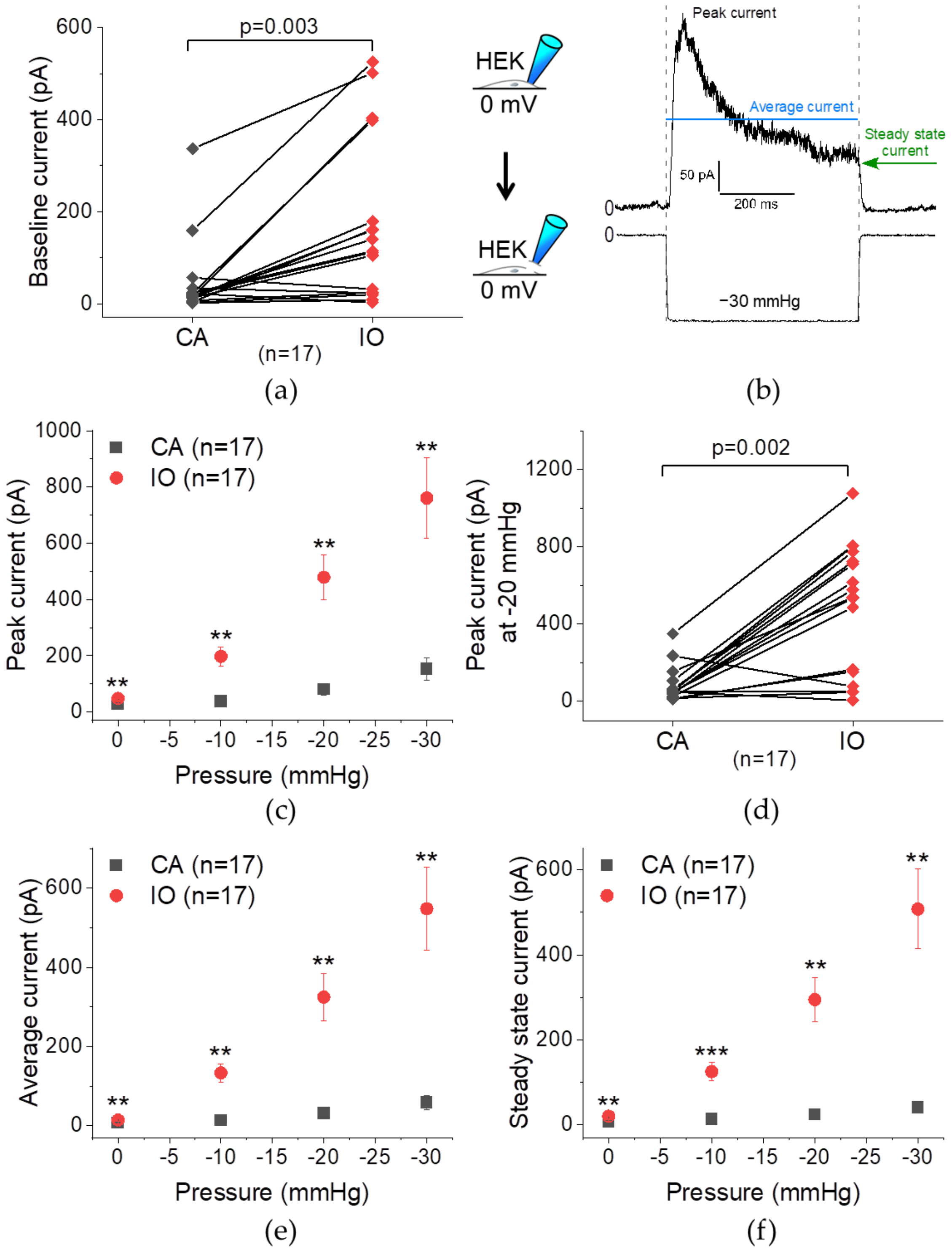


**Suppl. Figure 8.** Paired mechanically-induced TREK-1 activity in cell-attached (CA, grey) and inside-out (IO, red) configuration. TREK-1 was transfected at 0.1 µg/dish in HEK cells. Paired patch-clamp measurements at a holding potential of 0 mV, stretch applied from 0 to −30 mmHg, consecutively in both configurations, 20 s after seal formation (for CA) and 30 s after patch excision (for IO); N = 1 passage; Wilcoxon signed rank test. (a) Single data points of baseline current at 0 mmHg. (b) Analyzed parameters, obtained for one pulse of pressure (vertical dashed lines); from the recorded current trace, the peak (black), average (blue), and near steady-state (green) current amplitudes are deduced. (c, d) Peak current−pressure relationship and single data points at −20 mmHg. (e) Average current−pressure relationship (average current over the pulse of pressure); (f) Steady-state current−pressure relationship.

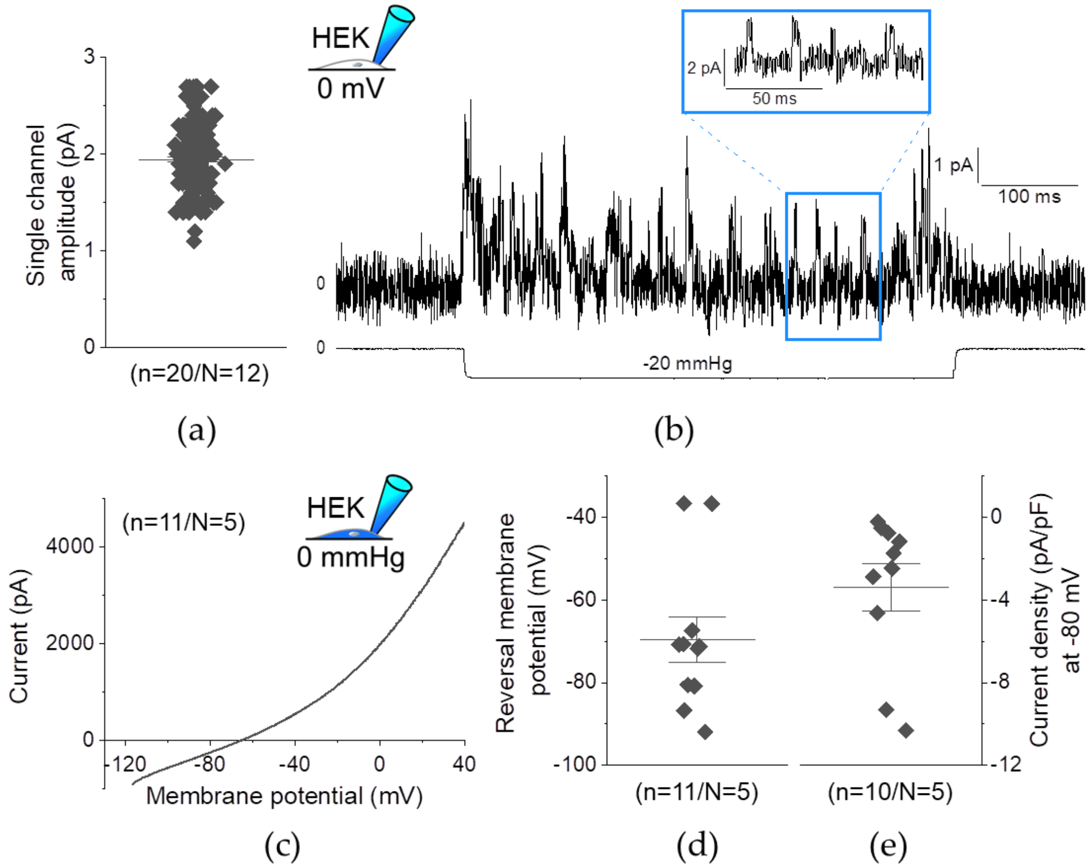


**Suppl. Figure 9.** Single channel amplitude and reversal membrane potential ($E_{rev}$) of TREK-1 (transfected at 0.1 µg/dish in HEK cells). Mechanically- and electrically-induced TREK-1 activity in cell-attached (holding potential 0 mV; a,b) and whole-cell configuration (pipette pressure 0 mmHg; c–e), respectively. (a) Single channel amplitudes, irrespective of the pressure pulse, multiple values/cell (in total, 185 values); average: −2.0 ± 0.0 pA. (b) Representative recording of single channel activity during a pressure pulse of −20 mmHg. (c) Current–membrane potential relationship, average of all recordings. (d) Single data points for $E_{rev}$; average: −69.5 ± 5.4 mV. (e) Single data points for current density at −80 mV. N = number of passages; n = number of cells.

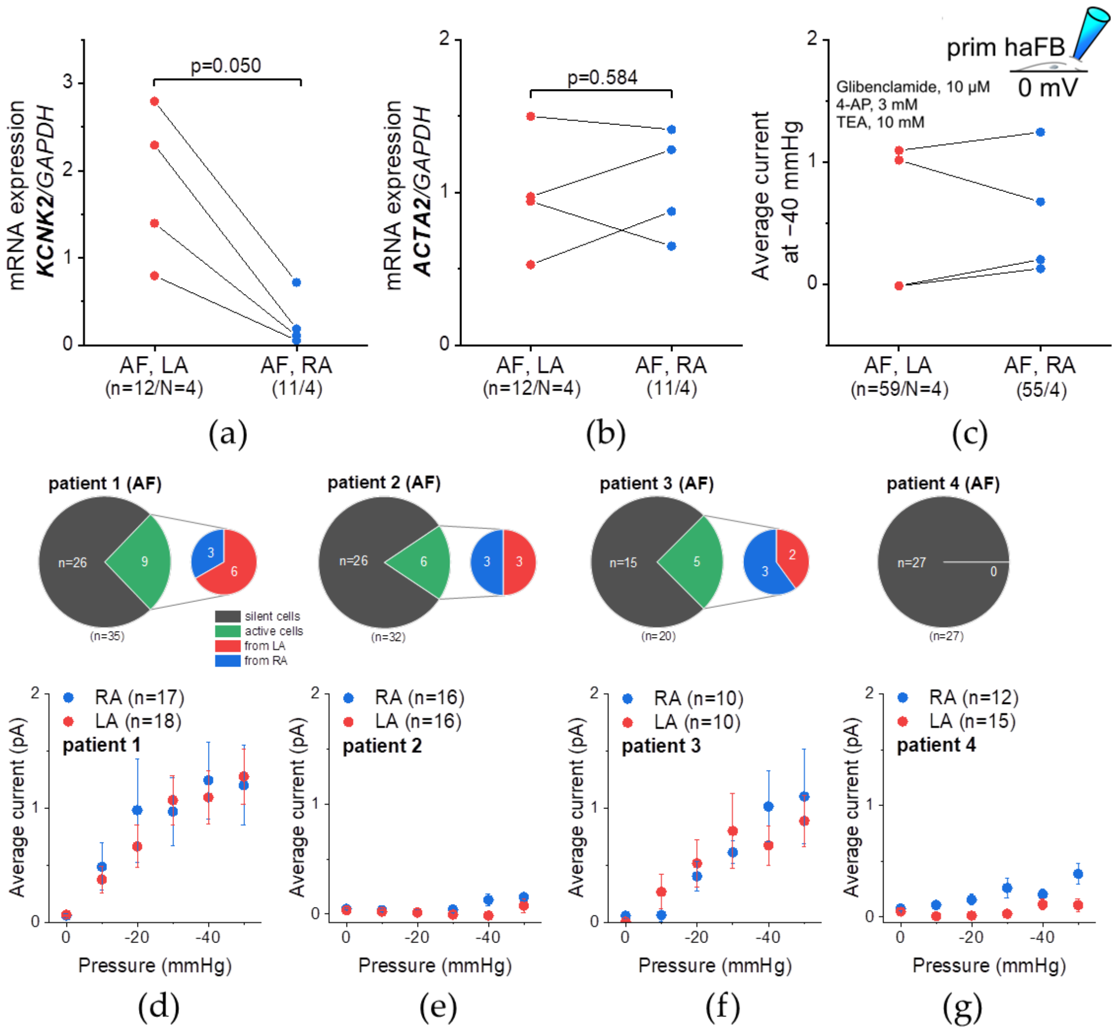


**Suppl. Figure 10.** Paired TREK-1 mRNA expression (a,b) and stretch-induced activity (c-g) in primary left (LA, red) and right (RA, blue) human atrial fibroblasts (HAFprim) from patients in atrial fibrillation (AF). (a, b) qPCR results of *KCNK2* (TREK-1; a) and *ACTA2* (αSMA; b) mRNA expression normalized to *GAPDH*; Wilcoxon signed rank test. (c−g) Patch-clamp measurements in inside-out configuration at a holding potential of 0 mV in the presence of non-$K_{2P}$ channel inhibitors (c, inset). (c) Average current at −40 mmHg; Wilcoxon signed rank test. (d−g, top row) Number of active cells (green) and silent cells (grey) for individual patients in AF; active cells are further divided into their tissue provenance. (d−g, bottom row) Average current (over the duration of the pressure pulse)−pressure relationship from 0 to −50 mmHg, comparing RA to LA for individual patients in AF; Mann−Whitney test. N = number of patients, n = number of dishes/cells.

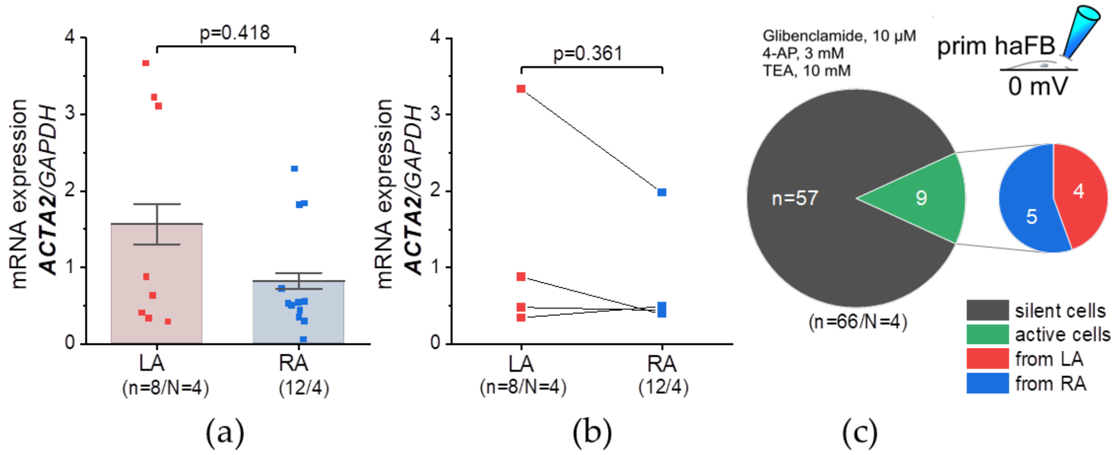


**Suppl. Figure 11.** Paired TREK-1 mRNA expression and stretch-induced activity in primary left (LA, red) and right (RA, blue) porcine atrial fibroblasts. (a, b) qPCR results of *KCNK2* (TREK-1; not detected) and *ACTA2* (αSMA) mRNA expression normalized to *GAPDH*; Mann–Whitney (a) and Wilcoxon signed rank (b) test. (c) Patch-clamp measurements in inside-out configuration at a holding potential of 0 mV in the presence of non-$K_{2P}$ channel inhibitors (c, inset); number of active cells (green) and silent cells (grey); active cells are further divided into their tissue provenance: LA (red) or RA (blue). N = number of pigs, n = number of dishes/cells.

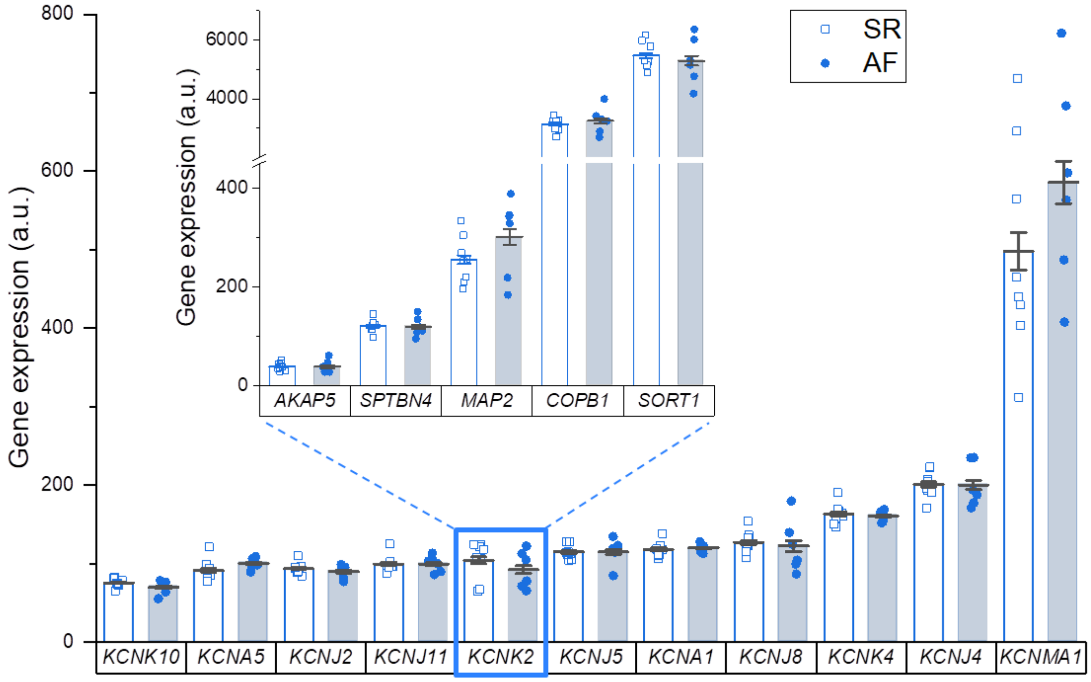


**Suppl. Figure 12.** Gene expression of $SAC_K$ and TREK-1 regulatory partners (inset) in primary human right atrial fibroblasts obtained from patients in sinus rhythm (SR, □) or sustained atrial fibrillation (AF, ●); assessed by gene micro-array; gene expression expressed in arbitrary units (a.u.); one data point represents one patient; SR: N = 8 patients; AF: N = 6 patients; genes are ordered with increasing abundance in SR; Mann–Whitney test. Results: $SAC_K$ gene expression in $HAF_{prim}$ derived from patients in SR ranged from 75.2 ± 2.0 for *KCNK10* (TREK-2) to 498.0 ± 135.7 for *KCNMA1*, with *KCNK2* being rather lowly expressed (103.9 ± 8.7). *SORT1* and *COPB1* were highly expressed (5470.0 ± 160.5 and 3133.1 ± 84.9, respectively) and *AKAP5*, *SPTBN4* and *MAP2* were lowly expressed (38.9 ±2.8, 120.9 ± 4.8 and 255.9 ± 16.8, respectively).

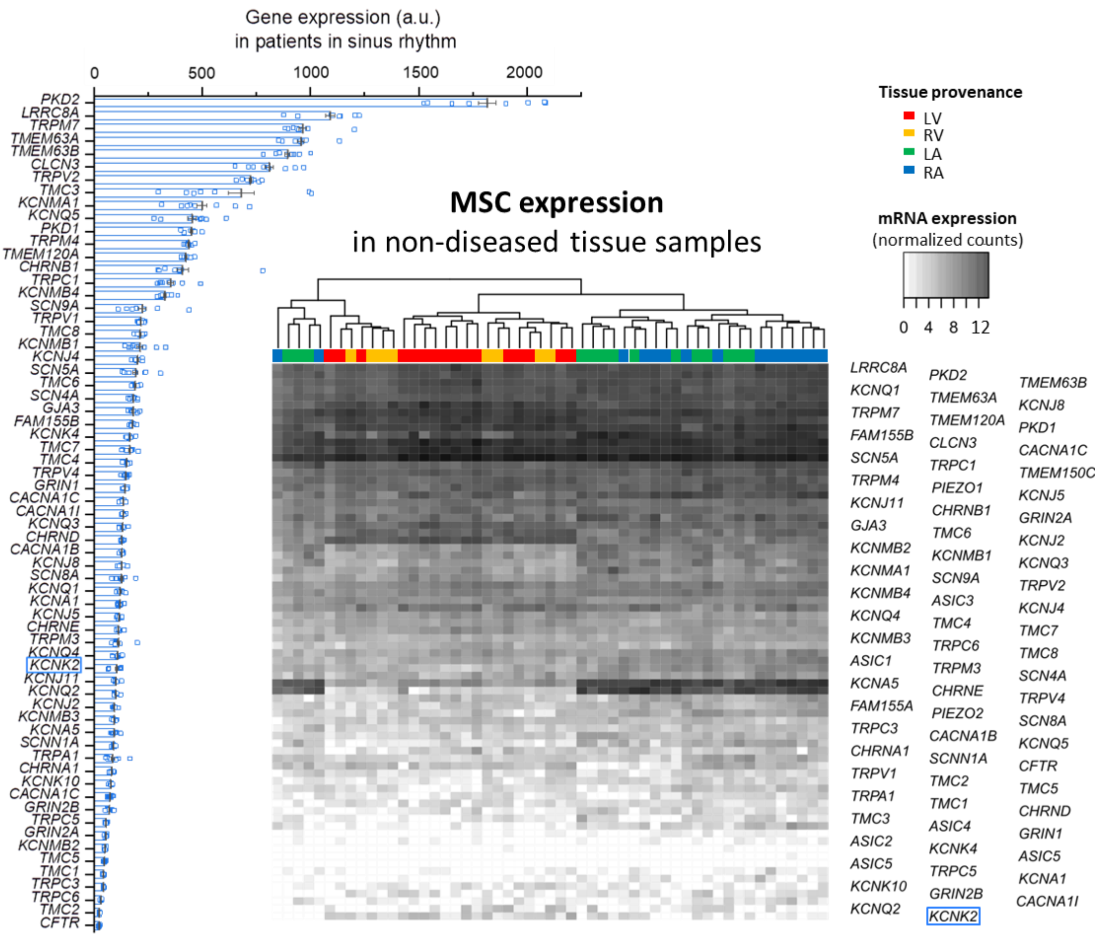


(a) (b)

**Suppl. Figure 13.** MSC mRNA expression in primary human right atrial (RA) fibroblasts obtained from patients in sinus rhythm (a) or in non-diseased human cardiac tissue samples (b); TREK-1 is highlighted in blue. (a) Gene expression assessed by gene micro-array; expressed in arbitrary units (a.u.); one data point represents one patient; N = 8 patients; genes are ordered with increasing abundance; no MSC was differentially expressed in AF (Mann−Whitney test). (b) Gene expression assessed by RNA-sequencing; expressed in normalized counts derived from the comparison atria *vs.* ventricles; tissue provenance: left ventricle (LV), right ventricle (RV), left atrium (LA) and RA; each row represents data from one tissue sample; each line represents one MSC gene.

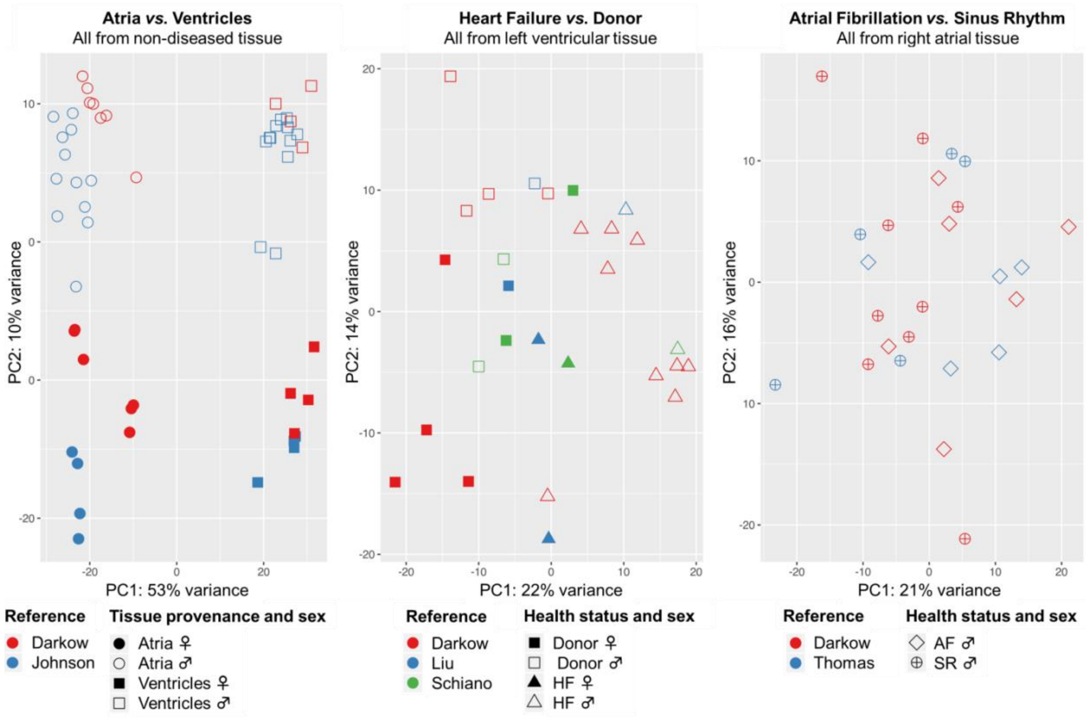


(a) (b) (c)

**Suppl. Figure 14.** Principal component (PC) analysis illustrating inter- and intra-group variability of samples *per* comparison: (a) atria (including left and right atrium) *vs.* ventricles (including left and right ventricle), in non-diseased (donor) tissue; (b) heart failure (HF; including dilated and ischemic cardiomyopathy) *vs.* donor, in left ventricular tissue; (c) atrial fibrillation (AF) *vs.* sinus rhythm (SR; including coronary artery and heart valve disease), in right atrial tissue. ♂: male patient/donor; ♀: female patient/donor; each individual point represents data from one tissue sample.

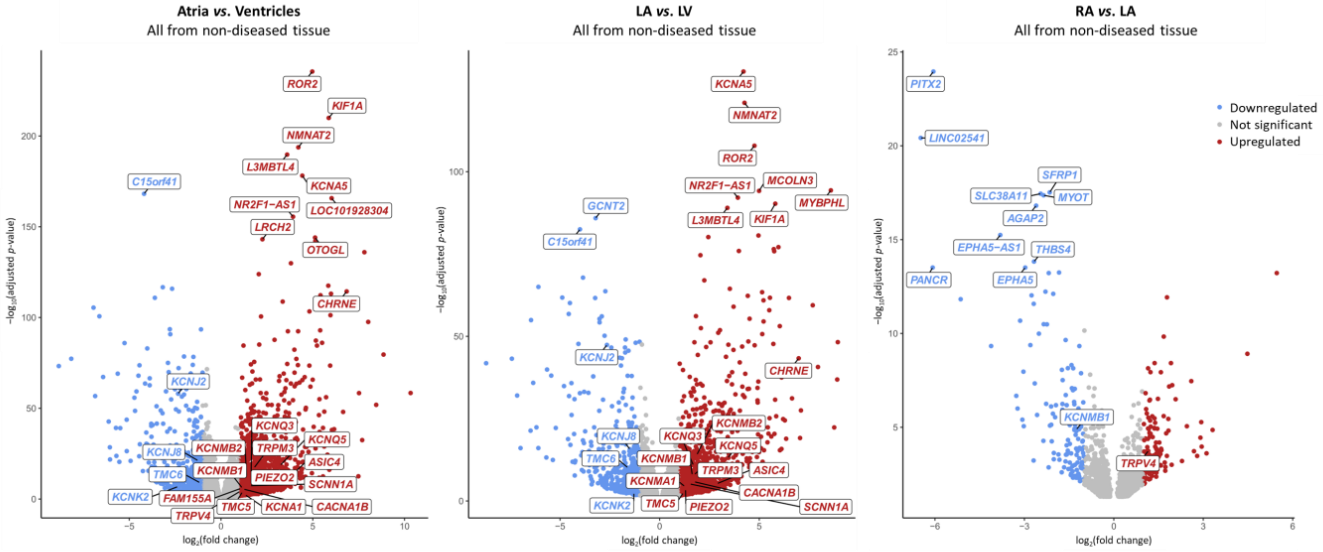


(a) (b) (c)

**Suppl. Figure 15.** Chamber-selective mRNA expression in non-diseased human cardiac tissue samples. Each dot represents one gene; among the genes with adjusted *p*-value < 0.05, MSC and top 10 differentially expressed genes are labelled. Genes significantly ($log_2$(fold change) > |1|) higher (red) or lower (blue) expressed (a) in the atria than in the ventricles; (b) in the left atrium (LA) than in the left ventricle (LV); (c) in the right atrium (RA) than in LA.

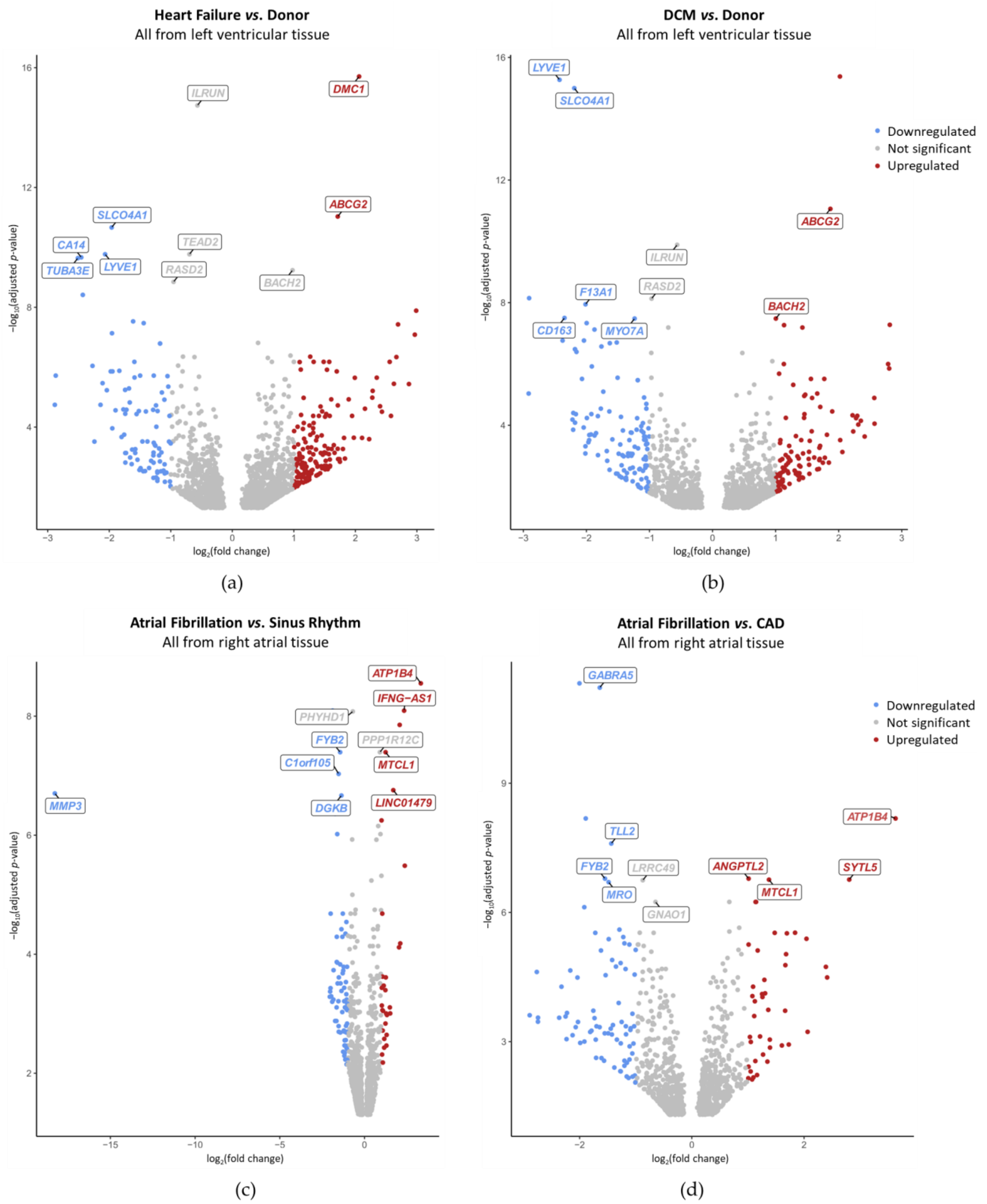


**Suppl. Figure 16**. Disease-selective mRNA expression in left ventricular (a, b) and right atrial (c, d) human cardiac tissue samples. Each dot represents one gene; among the genes with adjusted *p*-value < 0.05, MSC and top 10 differentially expressed genes are labelled. Genes significantly ($\log_2$(fold change) > |1|) higher (red) or lower (blue) expressed are colored. (a, b) Heart failure (HF; pooled from dilated cardiomyopathy [DCM] and ischemic cardiomyopathy; a) and DCM (b) compared to non-diseased donor hearts. (c, d) Atrial fibrillation (AF) compared to sinus rhythm (SR; pooled from coronary artery disease [CAD] and heart valve disease; c) and compared to CAD (d).

**Suppl. Table 1.** Human mechano-sensitive ion channels (MSC); alphabetical order. Cardiac MSC were mainly selected based on [126] and [4]. Additional candidate MSC were extracted from the gene ontology resource [127, 128] (GO:0008381; organism: homo sapiens) where they have been assigned mechano-sensitivity based on phylogenetic analysis [110] and sequence similarity [111]. When mechano-sensitivity has been attributed to specific subunits only these were selected, if not, all subunits were considered.

| Gene | Protein | Category | Reference |
|---|---|---|---|
| *ASIC1,2,3,4,5* | Acid-sensing ion channel 1,2,3,4,5 | MMC ($Na^+$) | [129] |
| *CACNA1B,C,I* | Voltage-dependent N,L,T-type $Ca^{2+}$ channel subunit $\alpha$-1B,C,I or $Ca_v$2.2, 1.2, 3.3 | MMC ($Ca^{2+}$) | [130], [131], [132] |
| *CFTR* | Cystic fibrosis transmembrane conductance receptor | MMC ($Cl^-$) | [133] |
| *CHRNA1,B1,D,E* | Acetylcholine receptor nicotinic subunit $\alpha$1,$\beta$1,$\delta$,$\varepsilon$ | MMC | [134] |
| *CLCN3* | $H^+$/$Cl^-$ exchange transporter 3 or CLC-3 | MMC ($Cl^-$) | [135] |
| *FAM155A,B* | Transmembrane protein FAM155A,B | candidate | [110] |
| *GJA3* | Gap junction $\alpha$-3 protein or Connexin 46 | MMC | [136] |
| *GRIN1, 2(A,B)* | Ionotropic glutamate receptor NMDA type subunit 1, 2 | | [137], [138] |
| *KCNA1,5* | $K^+$ voltage-gated channel subfamily A member 1,5 or $K_v$1.1,1.5 | $SAC_K$ | [139], [140] |
| *KCNJ2,4* | Inward rectifier $K^+$ channel 2,4, or $K_{ir}$2.1,2.3 | | [141], [142] |
| *KCNJ5* | G protein-activated inward rectifier $K^+$ channel 4 or $K_{ir}$3.4 or GIRK4 | | [143] |
| *KCNJ8,11* | ATP-sensitive inward rectifier $K^+$ channel 8,11 or $K_{ir}$6.1,6.2 | | [144] |
| *KCNK2,4,10* | $K^+$ channel subfamily K member 2,4,10 or TREK-1, TRAAK, TREK-2 | | [15], [145], [146] |
| *KCNMA1* | $Ca^{2+}$-activated $K^+$ channel subunit $\alpha$-1 or $K_{Ca}$1.1 or $BK_{Ca}$ $\alpha$ | $SAC_K$, MMC ($K^+$) | [125] |
| *KCNMB1,2,3,4* | $Ca^{2+}$-activated $K^+$ channel subunit $\beta$-1,2,3,4 or BK$\beta$1,2,3,4 | | |
| *KCNQ1,2,3,4,5* | $K^+$ voltage-gated channel subfamily KQT member 1,2,3,4,5 or $K_v$7.1,7.2,7.3,7.4,7.5 | VAC | [147], [148], [149],[150] |
| *LRRC8A* | Leucine rich repeat containing 8 VRAC subunit A or SWELL1 | | [151, 152] |
| *PIEZO1,2* | Piezo-type mechanosensitive ion channel component 1,2 | $SAC_{NS}$ | [153] |
| *PKD1,2* | Polycystin-1,2 or Transient receptor potential cation channel subfamily P member 1,2 or TRPP1,2 | MMC ($Ca^{2+}$) | [154], [155] |

| | | | |
|---|---|---|---|
| *SCN(4,5,8,9)A* | Na$^+$ channel protein type 4,5,8,9 subunit α or Na$_V$1.4,1.5,1.6,1.7 | SAC$_{NS}$, MMC (Na$^+$) | [156], [157], [158], [159] |
| *SCNN1A* | Na$^+$ channel epithelial 1 subunit α or ENaCα | SAC$_{NS}$ | [160] |
| *TMC1,2,3,4,5,6,7,8* | Transmembrane channel like protein 1,2,3,4,5,6,7,8 | MMC (Cl$^-$), candidate | [161], [110] |
| *TMEM120A* | TACAN | SAC$_{NS}$ | [162] |
| *TMEM150C* | Transmembrane protein 150C or Tentonin 3 | MMC | [163] |
| *TMEM63A,B* | CSC-1 like protein 1,2 | SAC$_{NS}$, candidate | [164], [111] |
| *TRPA1* | Transient receptor potential cation channel subfamily A member 1 | SAC$_{NS}$ | [165] |
| *TRPC1,3,5,6* | Short transient receptor potential channel 1,3,5,6 | | [166], [167], [168], [169] |
| *TRPM3,4,7* | Transient receptor potential cation channel subfamily M member 3,4,7 | | [170], [171], [172] |
| *TRPV1,2,4* | Transient receptor potential cation channel subfamily V member 1,2,4 | | [173], [174], [175] |

MMC, mechanically modulated ion channels; MGC, mechanically gated ion channels; VAC, volume-activated ion channels; SAC, stretch-activated ion channels; SAC$_K$, K$^+$-selective SAC; SAC$_{NS}$, cation non-selective SAC.

**Suppl. Table 2.** TREK-1 regulatory partners.

| Gene | Protein | Effect | Reference |
|---|---|---|---|
| *AKAP5* | A-kinase-anchoring protein AKAP150 | opening TREK-1 and making it insensitive to lipids, stretch and pH | [176] |
| *SPTBN4* | Actin-associated protein βIV-spectrin | TREK-1 membrane targeting | [177] |
| *MAP2* | Microtubule-associated protein Mtap2 | Enhancing TREK-1 surface expression and activity | [178] |
| *COPB1* | Vesicle transport protein β-COP | | [179] |
| *SORT1* | Neurotensin receptor 3 NTSR3 or gp95/sortilin | | [66] |

**Suppl. Table 3.** Summary of chamber-selective MSC gene expression in non-diseased human cardiac tissue samples. Genes significantly (adjusted *p*-value < 0.05 and log2(FC) > 1) higher expressed in the heart chamber indicated in the column than in the heart chamber indicated in the row; grey cells indicate that the corresponding comparisons could not be performed with our dataset; genes are sorted in alphabetical order.

| ↗ ↘ | **LV** | **RV** | **LA** | **RA** | |
|---|---|---|---|---|---|
| **LV** | | | *ASIC1,4*<br>*CACNA1B*<br>*CHRNE*<br>*KCNA5*<br>*KCNMA1* | | **Atrial-selective expression** |

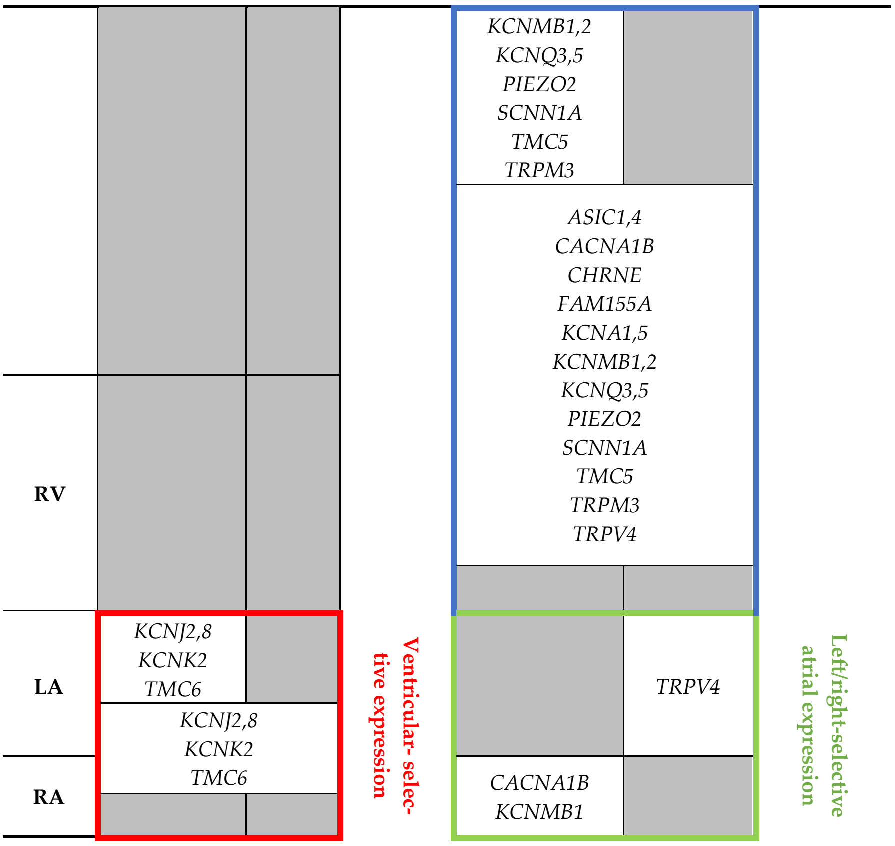


LV, left ventricle; RV, right ventricle; LA, left atrium; RA, right atrium.

**Suppl. Table 4.** Summary of chamber-selective differential MSC expression in non-diseased human cardiac tissue samples. Upregulation (↑), downregulation (↓), or no significant difference (−) are indicated such that the symbols outside parentheses sum up our experimental observations, and the symbol within parentheses previously reported findings from the literature. Background color indicates that our observations match (green) or do not match (orange) prior reports; alphabetical order.

| Gene | Atria *vs.* Ventricles | LA *vs.* LV | RA *vs.* LA |
|---|---|---|---|
| *ASIC1* | ↑ | ↑ | – |
| *ASIC4* | ↑ | ↑ | – |
| *CACNA1B* | ↑ | ↑ | ↓ |
| *CHRNE* | ↑ | ↑ | – |
| *FAM155A* | ↑ | – | – |
| *KCNA1* | ↑ | – | – |
| *KCNA5* | ↑ | ↑ (↑ [104, 105]) | – |
| *KCNJ2* | ↓ | ↓ (↓ [104, 105]) | – |
| *KCNJ8* | ↓ | ↓ | – |
| *KCNK2* | ↓ | ↓ | – |
| *KCNMA1* | – | ↑ | – |
| *KCNMB1* | ↑ | ↑ | ↓ |
| *KCNMB2* | ↑ | ↑ | – |
| *KCNQ3* | ↑ | ↑ | – |
| *KCNQ5* | ↑ | ↑ | – |
| *PIEZO2* | ↑ | ↑ | – |
| *SCNN1A* | ↑ | ↑ | – |
| *TMC5* | ↑ | ↑ | – |
| *TMC6* | ↓ | ↓ | – |

| | | | |
|---|---|---|---|
| *TRPM3* | ↑ | ↑ | − |
| *TRPV4* | ↑ | − | ↑ |

LA, left atrium; LV, left ventricle; RA, right atrium; LA, left atrium.

**Suppl. Table 5.** Summary of disease-selective differential MSC expression. Upregulation (↑), downregulation (↓), or no significant difference (−) are indicated such that the symbols outside parentheses sum up our experimental observations, and the symbol within parentheses previously reported findings from the literature. Background color indicates that our observations match (green) or do not match (orange) prior reports; alphabetical order.

| Gene | HF *vs.* Donor | DCM *vs.* Donor | AF *vs.* SR | AF *vs.* CAD |
|---|---|---|---|---|
| *ASIC3* | ↓ | ↓ | − | − |
| *CFTR* | ↓ (↓ [106, 107]) | − | − | − |
| *CHRNE* | ↑ | − | − | − |
| *KCNJ4* | ↑ | − | ↑ | − |
| *KCNJ5* | − | − | ↓ | ↓ |
| *KCNJ11* | ↓ (↓ [108]) | − | − | − |
| *KCNQ4* | − | − | ↑ | ↑ |
| *KCNQ5* | − | − | ↓ | ↓ |
| *LRRC8A* | ↓ | ↓ | − | − |
| *PKD1* | − | − | ↑ | ↑ |
| *SCN9A* | ↓ | ↓ | − | − |
| *TMC5* | − | − | ↓ | ↓ |
| *TMEM120A* | − | − | − | ↑ |
| *TMEM63B* | ↓ | − | − | − |
| *TRPC6* | ↑ | ↑ (↑ [12]) | − | − |

HF, heart failure; DCM, dilative cardiomyopathy; AF, (sustained) atrial fibrillation; SR, sinus rhythm; CAD, coronary artery disease.
